# Using seasonal genomic changes to understand historical adaptation: parallel selection on stickleback in highly-variable estuaries

**DOI:** 10.1101/2020.11.24.396291

**Authors:** Alan Garcia-Elfring, Antoine Paccard, Timothy J. Thurman, Ben A. Wasserman, Eric P. Palkovacs, Andrew P. Hendry, Rowan D. H. Barrett

## Abstract

Parallel evolution is considered strong evidence for natural selection. However, few studies have investigated the process of parallel selection as it plays out in real time. The common approach is to study historical signatures of selection in populations already well adapted to different environments. Here, to document selection in action under natural conditions, we study six populations of threespine stickleback (*Gasterosteus aculeatus*) inhabiting bar-built estuaries that undergo seasonal cycles of environmental changes. Estuaries are periodically isolated from the ocean due to sandbar formation during dry summer months, with concurrent environmental shifts that resemble the long-term changes associated with postglacial colonization of freshwater habitats by marine populations. We used pooled whole-genome sequencing (Pool-WGS) to track seasonal allele frequency changes in these populations and search for signatures of natural selection. We found consistent changes in allele frequency across estuaries, suggesting a potential role for parallel selection. Functional enrichment among candidate genes included transmembrane ion transport and calcium binding, which are important for osmoregulation and ion balance. The genomic changes that occur in threespine stickleback from bar-built estuaries could provide a glimpse into the early stages of adaptation that have occurred in many historical marine to freshwater transitions.

## Introduction

Knowledge of the genomic targets of natural selection is central to understanding the mechanisms responsible for adaptive evolution at the molecular level. Generating this knowledge often involves comparing patterns of genomic differentiation (e.g., F_ST_) between populations adapted to distinct ecological conditions (e.g. Hoekstra et al. 2006; llardo et al. 2018). In such studies, natural selection is considered a strong candidate for the mechanism driving phenotypic diversification when multiple closely related but independently-evolved populations use the same genetic pathways to reach a shared adaptive solution to an environmental challenge. Here, we refer to this phenomenon as parallel evolution (Elmer and Meyer 2011). In vertebrates, studies of parallel evolution have provided insights into the genetic mechanisms underlying adaptation to freshwater (Colosimo et al. 2005; Barrett et al. 2008; Schluter et al. 2010; Hohenlohe et al. 2010; Kitano et al. 2010; Lescak et al. 2015; Rudman et al. 2019; Fang et al. 2020), novel pathogens (Alves et al. 2019), low oxygen availability (McCracken et al. 2009; Foll et al. 2014; Wang et al. 2014; Graham and McCracken 2019; Lim et al. 2019), crypsis (Comeault et al. 2016; Jones et al. 2018; Barrett et al. 2019), nutrient-limited environments (Riddle et al. 2018), and dissolved ion (H^+^) profiles (Haenel et al. 2019). However, nearly all studies of parallel evolution are retrospective in the sense that they investigate reasonably well adapted populations that are no longer experiencing the strong selection associated with an environmental change.

Retrospective approaches thus have difficulty detecting the specific genetic changes that were under natural selection during the initial habitat shift amongst the noise from local effects and stochastic processes that accumulates afterward (Elmer and Meyer 2011). A valuable addition to the inferential toolbox, then, is to study natural selection that takes place *during* parallel habitat shifts. Most studies adopting this selection-based approach have used artificial perturbations of genotypes or environments, which have revealed genomic targets of strong selection (e.g. Anderson et al. 2014; Soria-Carrasco et al. 2014; Nosil et al. 2018; Barrett et al. 2019). However, the manipulations involved in such tests leaves uncertain the extent to which similar selection would attend natural habitat shifts. One solution is to take advantage of serendipitous occurrences, such as extreme weather events like droughts (Grant et al. 2017), hurricanes (Donihue et al. 2018), heat waves (Coleman et al. 2020; Gurgel et al. 2020) and cold snaps (Campbell-Staton et al. 2017; Card et al. 2018). However, the location and timing of extreme events like these are unpredictable by nature, limiting the study of how natural populations respond to such events. An alternative strategy, then, might be to study selection as it occurs in natural populations experiencing large, yet predictable, shifts between alternative environments, like those that occur during seasonal changes (e.g. Behrman et al. 2018; Tourneur et al. 2020). Here, we search for the signature of natural selection in a set of populations that experience parallel seasonal changes in local conditions that likely resemble the early phase of a classic habitat transition. Specifically, we study allele frequency changes in stickleback populations in environments that alternate between marine-like (brackish) and freshwater conditions.

### Study system

The threespine stickleback (*Gasterosteus aculeatus*, ‘stickleback’ hereafter) is a classic model system for studying parallel evolution (Boughman et al. 2005; Colosimo et al. 2005; Jones et al. 2012a, 2012b; Deagle et al. 2013; Hendry et al. 2009, 2013; Lescak et al. 2015; Paccard et al. 2018; Haenel et al. 2019; Smith et al. 2020). Over the past approximately 12,000 years, marine stickleback have repeatedly colonized and become adapted to freshwater environments, often through parallel phenotypic changes (Reimchen 1983; Colosimo et al. 2005) linked to predator defence (Reimchen 2000; Marchinko 2009; Miller et al. 2019; Wasserman et al. 2020) and ion regulation (Gibbons et al. 2016, 2017; Hasan et al. 2017). The genomic basis of such adaptation is partly known. For instance, researchers have identified genes of large effect underlying differences in the number of bony armour plates (e.g. *Eda* gene, e.g. Colosimo et al. 2005), pelvic spines (e.g. *Pitx1* gene, Chan et al. 2010), and the ability to osmoregulate (e.g. *Kcnh4* gene, Jones et al. 2012a; *Atp1a1a* gene, McCairns and Bernatchez 2010; Jones et al. 2012b; Hasan et al. 2017). These genes have been found primarily through a retrospective approach of studying signatures of selection millennia after the initial colonization.

Stickleback populations in bar-built estuaries along the coast of California represent a natural system for studying parallel selection over seasonal timescales. These populations experience repeated bouts of strong and abrupt temporal changes driven by wet winters and dry summers. With heavy winter rains, increased water flow breaches the wall of sediment (i.e., ‘sandbar’) that, during the summer, typically isolates estuaries from the ocean. When rains subside, sandbars are re-built by wave action and sand deposition in the spring or summer, isolating estuaries from the ocean and creating coastal lagoons (Behrens et al. 2009; Behrens et al. 2013; Rich and Keller 2013). The changes in precipitation that lead to the build-up of sandbars and the subsequent breaching can result in drastic environmental shifts in, for example, predator abundance (Becker and Reining 2008), salinity (Williams 2014), and habitat structure (Heady et al. 2015). For example, in between breaching events when the estuaries are closed to the ocean, salinity levels diminish as a surface freshwater layer progressively increases in thickness (Williams 2014) and a shift takes place from lotic (i.e. moving) brackish water to lentic (i.e. pond-like) freshwater (Heady et al. 2015; Des Roches et al. 2020). These seasonal habitat shifts may be analogous to the environmental changes experienced by stickleback populations during postglacial marine-to-freshwater colonization events and are replicated both spatially (in different estuaries) and temporally (with seasonal changes in precipitation).

To study natural selection in action, we sampled stickleback from six bar-built estuaries at two time points between breaching events, when the estuaries were isolated from the ocean. Using a whole-genome SNP dataset, we characterized the extent of allele frequency change between the sampling times, which should reflect, at least in part, natural selection taking place during that interval. Then, to evaluate evidence of parallel natural selection, we ask several questions. (1) What genomic regions show relatively large changes in allele frequency across time in multiple estuaries? (2) For these regions, do the changes in allele frequency occur in parallel across estuaries? (3) Do genes putatively under parallel selection show enrichment of genetic functions consistent with the changes in the environment? By obtaining this information over a seasonal timescale in multiple estuaries, we hope to gain insight into the genetic changes driven by selection that may have occurred when postglacial stickleback populations first colonized freshwater environments from the ocean.

## Methods

### Field sampling and DNA extraction

In 2016, we sampled stickleback from six bar-built estuaries (Figure 1), three from small coastal watersheds (< 7 km^2^; Old Dairy, Lombardi, and Younger) and three that are relatively large watersheds (> 22 km^2^; Laguna, Scott, and Waddell; see Table S1 for full estuary names and size metrics). We sampled in the spring at the end of the breaching season upon completion of the sandbar (after winter rain), and again in the fall before the breaching season (before the winter rain, Figure S1). Thus, we are testing for selection during the part of the year that estuaries are isolated from the ocean, which provides a single-season analogue of the marine to freshwater transition that is a classic theme in stickleback research (e.g. Colosimo et al. 2005; Bassham et al. 2018; reviewed by Hohenlohe and Magalhaes 2019). At each time point in each estuary, we collected 40 adult stickleback (> 30 mm in length) by means of minnow traps and beach seines. Although sampling time is less than one generation (~6 months), some stickleback may have given birth shortly after our first sample, with progeny growing large enough to be sampled as adults in our second sample. Selection during our sampling period therefore reflects both differential mortality and reproduction. Fish were euthanized with tricaine methanosulphonate (MS-222) and tissue samples (pectoral fin) were stored in 95% ethanol prior to DNA extraction. Collections were made in accordance with California Scientific Collector’s Permit SC-12752. Animal handling protocols were approved by the University of California, Santa Cruz IACUC under protocols Palke-1306 and Palke-1310. We extracted DNA following a standard phenol-chloroform procedure. Briefly, tissue samples were placed in digestion buffer containing proteinase K and incubated at 55 °C. We then isolated DNA using an isoamyl-phenol-chloroform solution, followed with ethanol precipitation.

**Figure 1.**
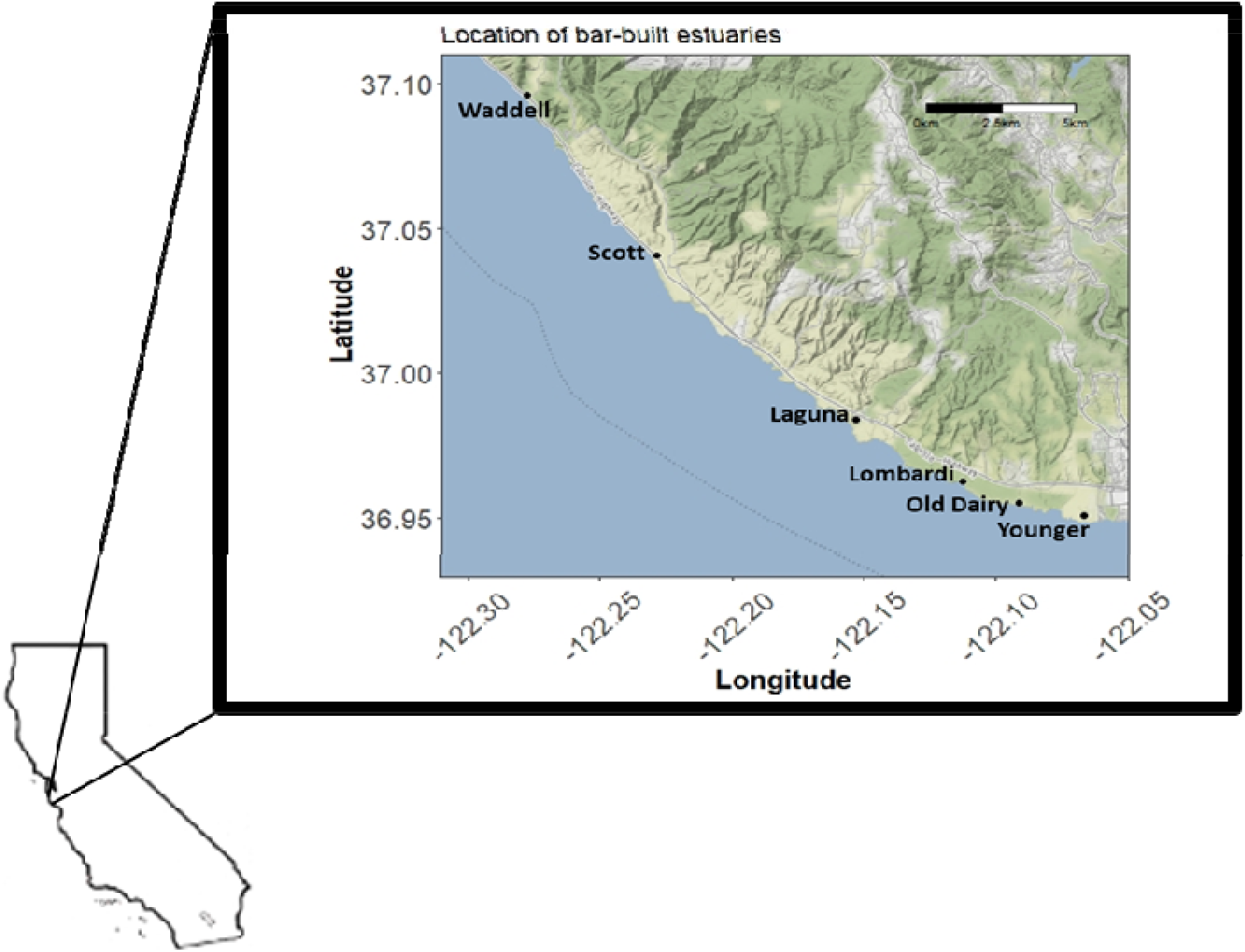
Locations of six bar-built estuaries sampled along the coast of California USA.

### Sequencing

We quantified all samples using a Picogreen^®^ ds DNA assay (Thermo Fisher Scientific, Waltham, USA) on an Infinite^®^ 200 Nanoquant (Tecan Group Ltd. Männedorf, Switzerland). Samples were normalized to a dsDNA concentration of 15ng/μl, re-quantified, and pooled according to sampling location and time of sampling. Thus, we created 12 pools of 40 individuals each (i.e. six estuaries sampled at two time points). Whole-genome libraries of each pool were prepared at the McGill University and Genome Quebec Innovation Center, Montreal, Canada, and sequenced across five lanes of Illumina HiSeq2500 with paired-end, 125bp reads.

### Bioinformatics

We filtered raw reads based on quality (--quality-threshold 20) and length (--min-length 50) with the *trim-fastq.pl* script of *Popoolation* (Kofler et al. 2011a). The resulting reads were mapped to the stickleback reference genome (BROADS S1) using *BWA mem* v. 0.7.13 (Li and Durbin 2009). We then used *SAMtools* (Li et al. 2009) to convert SAM files to BAM format and remove reads with mapping quality below 20 (samtools view -q 20). We then generated a mpileup file (samtools mpilep -B) and filtered for a minimum depth of coverage of 5X. We converted the mpileup file to the synchronized (sync) format using *Popoolation2* (Kofler et al. 2011b) for downstream analysis.

### Analysis of repeated genomic differentiation

In this study we use F_ST_ to measure changes in allele frequency within a lineage across time points (rather than differentiation between two lineages, Burri 2017). First, within each estuary, we calculated F_ST_ at the SNP level (Hartl and Clark 1997) to identify variants showing relatively large changes in allele frequency (i.e. outliers) between the two time points. We then quantified the extent of overlap in these outliers among estuaries. To obtain genome-wide F_ST_ estimates, we applied the *fst-sliding.pl* script of *Popoolation2* for each of the six estuaries post-breaching (i.e. brackish conditions) versus pre-breaching (freshwater) (--min-count 2, --min-coverage 5, --max-coverage 100, --min-covered-fraction 0, --window-size 1, --step-size 1, --pool-size 40:40:40:40:40:40:40:40:40:40:40:40, --suppress-noninformative). We only analyzed genomic regions assembled at the chromosome level (i.e. scaffolds excluded). We included data from chromosome 19 (the allosome) as we did not find evidence of any mapping bias (mean = 23.92, SD: 6.48, range 5-74) on this chromosome relative to the genome-wide average, or any strong signal of consistent allele frequency change (see Results). We identified F_ST_ outliers as SNPS that fell in the top 5% of the F_ST_ distribution. These loci were excluded from calculations of genome-wide F_ST_ and allele frequency change distributions to obtain estimates for putatively neutral SNPs (e.g. Batista et al. 2016). To discover candidates potentially under selection, we focused on SNPs that showed large allele frequency changes in multiple estuaries. Because drift and sampling variance will affect loci at random across the genome within any particular estuary, it is unlikely that consistent genetic changes across three or more different estuaries will be due to stochastic processes. We used a custom bash script to quantify F_ST_ outlier overlap across estuaries and identify SNPs that qualify as outliers in at least three out of the six estuaries. However, because evidence of repeated changes in allele frequency in the same SNP (as shown by F_ST_) does not necessarily mean that these changes were parallel (i.e. in estuary *X* an allele shows a large *increase* in frequency, while in estuary *Y* the same allele experiences a large *decrease* in frequency), we also tested for parallelism in allele frequency change.

### Parallel changes in allele frequencies in response to seasonality

We identified SNPs showing consistent directional changes in allele frequency across our estuaries using the program *PoolFreqDiff* (Wiberg et al. 2017). *PoolFreqDiff uses* a generalized linear model with a quasibinomial error distribution (qGLM). Wiberg et al. (2017) showed that the qGLM has a substantially lower false positive rate than the Cochran-Mantel-Haenszel test, a method commonly used in pool-seq studies to identify consistent changes in allele frequency across replicates. We used the same flags (e.g. minimum read count and coverage settings) in the *PoolFreqDiff* program as in our F_ST_ analysis. The qGLM test implemented in *PoolFreqDiff* has the potential to identify parallel selection reflected as subtle, but consistent, changes in allele frequency in multiple estuaries. Such small changes in allele frequency are unlikely to be identified as F_ST_ outliers in individual estuaries. We used the ‘no rescaling’ option of *PoolFreqDiff* (re-scaling allele counts relative to the effective sample size gave similar results). We corrected for population structure using the empirical null-hypothesis approach (Caye et al. 2016; François et al. 2016). Visual inspection of the histogram of corrected P-values confirmed a uniform distribution under the null-hypothesis, indicating that confounders were controlled (Figure S2). Next, we corrected for multiple hypothesis testing using the false discovery rate (FDR) procedure implemented in the R package *qvalue* V2.14 (Storey et al. 2018). We analyzed three sets of outliers to study potential targets of selection. First, to look for strong and parallel changes in allele frequency, we categorized ‘F_ST_-qGLM outliers’ as SNPs that are an F_ST_ outlier in at least three estuaries and also significant (FDR = 1%) under the qGLM model. Second, we identified outliers from each of the two distinct approaches (F_ST_ and qGLM) but with more conservative thresholds than those used in the overlapping F_ST_-qGLM outlier set. For ‘F_ST_ candidates’, we identified F_ST_ outliers (SNPs in the top 5% of the F_ST_ distribution in a single estuary) that were shared across at least four of six estuaries (as opposed to the three estuaries minimum requirement in the F_ST_-qGLM outlier set). Note that the frequency changes across these estuaries may not be parallel and thus this outlier set accounts for potential causes of selection that may differ in direction among estuaries. We also tested whether estuary size may influence the likelihood of shared targets of selection (F_ST_ outliers; see supplemental information). Finally, ‘qGLM outliers’ are SNPs identified as highly significant using the quasibinomial GLM test for parallel changes in allele frequency, here using an FDR = 0.01% as opposed to the less conservative FDR = 1% used for the F_ST_-qGLM outlier set. We obtained estimates of allele frequency change across time points for F_ST_-qGLM outliers and putatively neutral loci with respect to F_ST_ (not in top 5% F_ST_ distribution) using the *snp-frequency-diff.pl* script of *Popoolation2*.

### Identification of candidate genes and analysis of molecular function

To identify genes putatively under parallel selection, we used a custom bash script to map outliers (i.e. F_ST_-qGLM outliers, F_ST_ candidates, and qGLM outliers) to protein-coding genes in the reference genome. We limited our search to a set of 14,252 protein-coding gene annotations with attributes “ID=gene” and “biotype=protein_coding”. To gain insights into the traits under selection, we analyzed candidate genes for enrichment of molecular functions. To do this, we obtained gene names and gene ontology (GO) information from the stickleback reference genome on *Ensembl* using the R package *biomaRt* (Smedley et al. 2009). We compared the three lists of candidate genes with the reference set of 14,252 genes (‘gene universe’) and tested for functional enrichment using the package *TopGO* 2.34.0 (Rahnenfuhrer 2018) and the Fisher’s exact test. To reduce false positives, we pruned the GO hierarchy by requiring that each GO term have at least 10 annotated genes in our reference list (“nodeSize = 10”).

## Results

### Data processing and F_ST_ estimates in response to seasonal sandbar formation

Our sequencing efforts led to an average of 23,914,973,875 bases sequenced per pool (SD: 1,760,685,042). After filtering data, we obtained 101,911,501 bases for variable sites, providing F_ST_ estimates for 4,024,542 SNPs distributed across 21 stickleback chromosomes. The average minimum coverage per SNP was 25.32 (SD = 6.96, range: 5 – 84, Figure S3) among pools. Overall, allele frequency changed relatively little within estuaries, showing a mean ‘neutral’ F_ST_ of 0.0253 across time points (Waddell = 0.0224; Lombardi = 0.0230; Old Dairy = 0.0216; Younger = 0.0243; Scott = 0.0236; Laguna = 0.0369).

### Consistent changes in allele frequency and the signature of parallel selection

To identify candidates under temporally varying parallel selection, we looked for an overlap among estuaries of SNPs that fall in the top 5% of the F_ST_ distribution. As expected, most SNPs found in the top 5% of the distribution in an estuary only reach this threshold in a single estuary (Figure 2). Yet, we find 22,111 SNPs in the top 5% in three or more estuaries. The majority of these SNPs, 19,390 SNPs (87.7%), are confined to exactly three estuaries, with 2,721 ‘F_ST_ candidates’ found in four or more estuaries. At a FDR of 1%, we identified 37,687 SNPs using *PoolFreqDiff*, 705 of which overlapped with the SNPs that were F_ST_ outliers in at least three estuaries (‘F_ST_-qGLM outliers’; Figure 3 and Figure 4). We also identified 2,411 SNPs with a FDR of 0.01% (‘qGLM outliers’).

**Figure 2.**
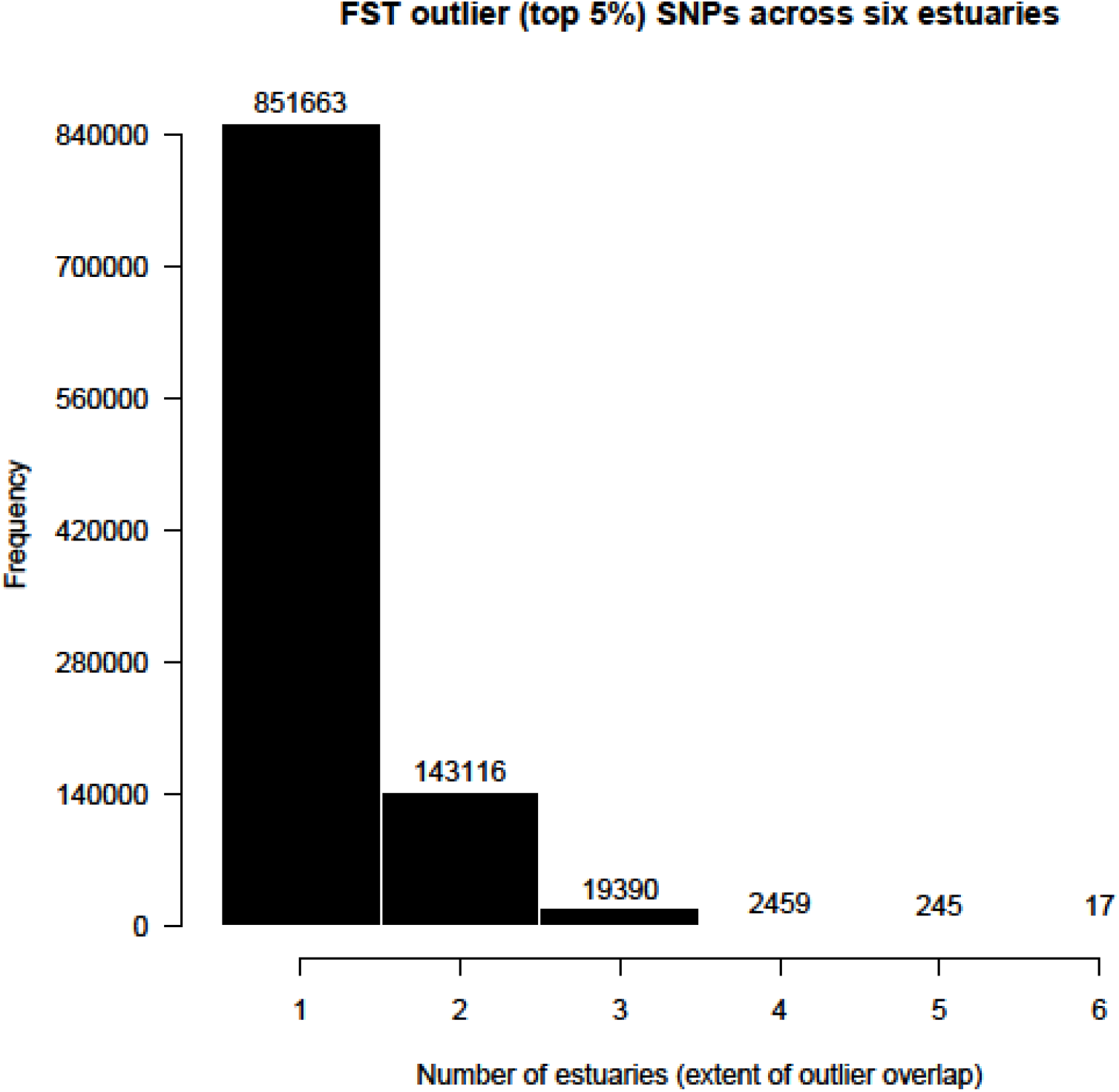
Extent of overlap of F_ST_ outlier loci across six estuaries.

**Figure 3.**
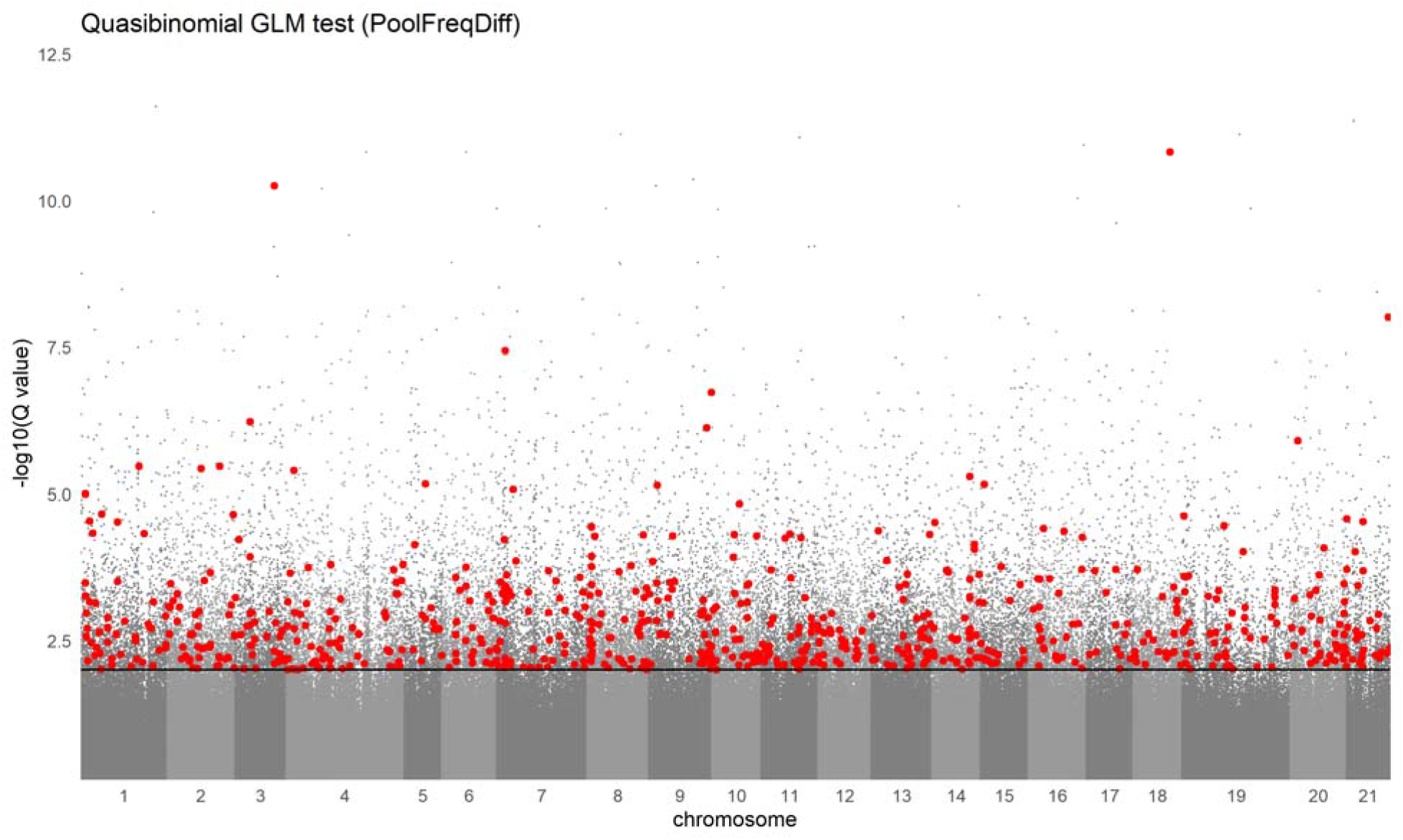
Result of qGLM test (*PoolFreqDiff*) for parallel changes in allele frequency. Each dot represents a single SNP (MT chromosome and unplaced scaffolds excluded). The 705 loci identified by F_ST_ analysis as candidates in at least three estuaries and by the qGLM test as significant (FDR = 1%) are shown in red (‘F_ST_-qGLM outliers’). Black line demarks the 1% false discovery rate.

**Figure 4.**
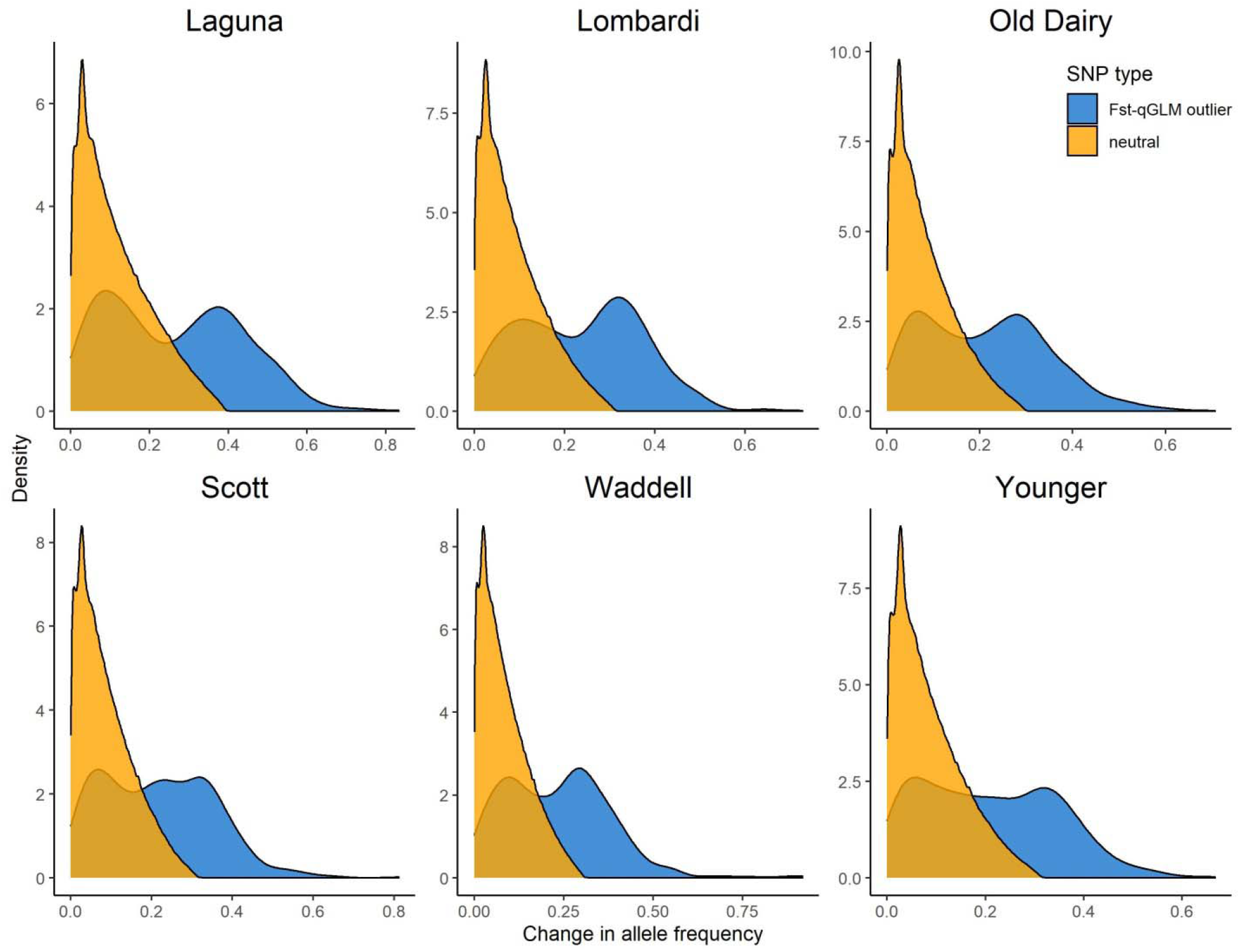
Distribution of allele frequency change in genome-wide SNPs and F_ST_-qGLM outliers.

### Candidate genes and analysis of molecular function

We mapped outlier SNPs to genes and found 710 genic F_ST_ candidates in 579 different genes (Table S3), 704 qGLM outliers in 569 genes (Table S4), and 190 F_ST_-qGLM outliers in 169 different genes (Table S5). All three sets of outliers have candidate genes associated with ion transfer, including *Wnk4* (Table S3 and Table S4) and *Nalcn* (Table S4 and Table S5). We find consistent changes in allele frequency in ATPase genes that code for proteins that transport, for example, sodium and potassium (e.g. *Atp1b1a*, Table S3) and phospholipids (*Atp8b5a*, Table S3 and Table S5). Potassium transport channels are also found among our candidate genes (e.g. *Kcnma1a*, Table S4 and Table S5; *Kcnn1a*, Table S3) as well as genes involved in calcium binding or transport, like the calcitonin receptor (*Calcr*, Table S3 and Table S4), calmodulin (*Calm1b*, Table S3) and the calcium channel *Cacna1d* (Table S5). Yet others code for various mitogen-activated protein kinases (e.g. *Map3k12*, Table S4; *Mapkbp1*, Table S5). We also found an F_ST_ outlier mapping to *Ccny*, a gene involved in the cell cycle (Table S3).

### GO analysis

For an overall assessment of the gene functions that are most represented among our three sets of candidate genes, we tested for enrichment of molecular function. We find that the candidate genes from all three outlier sets have in their top ten most significant GO terms molecular functions related to ion channel activity. For example, F_ST_ candidates show enriched GO functions related to calcium ion binding (Table S6). Among the most enriched molecular functions among qGLM candidate genes are metal ion transmembrane transporter activity and calcium ion transmembrane transporter activity (Table S7). Similarly, F_ST_-qGLM outlier genes are enriched for ion transmembrane transporter activity and ligand-gated ion channel activity (Table S8). We also found extracellular matrix structural constituent (Table S6) among the significant molecular functions (see Discussion).

## Discussion

To document natural selection, we studied stickleback populations from estuaries experiencing seasonal fluctuations in environmental conditions. We found evidence of parallel selection on genes linked to ion transport and salinity adaptation. Consistent with a change in the ionic environment, we found that the most statistically significant functions were related to ionic homeostasis. Our findings suggest that intermittent connectivity with the ocean results in episodic shifts in selection regime, a change that may resemble the initial phase of freshwater colonization that occurred during the marine-freshwater transitions of postglacial stickleback populations. More generally, our study adds to a growing literature that collectively shows that natural selection can drive genetic change over very short timespans.

### Parallel selection over a seasonal timescale

We found evidence of natural selection in the form of consistent changes in allele frequency in stickleback populations from different bar-built estuaries. Our results showed changes in allele frequency on genes with functions related to ion balance. This suggests that seasonal fluctuations of environmental conditions shift the selection regime within bar-built estuaries. The time scale involved in the overall subtle but consistent changes in allele frequency detected here conforms with studies that show selection for freshwater adaptation can be detected within a single year rather to decades (Bell 2001; Kristjánsson et al. 2002; Bell et al. 2004; Lescak et al. 2015; Marques et al 2018). These results are also consistent with the idea that adaptation to a particular environment, like freshwater, likely happens in the first few generations after colonization (e.g. reviewed by Reimchen et al. 2013). Importantly, our results set a new bar for how quickly selection can result in genetic adaptation during freshwater colonization by stickleback.

### Seasonal allele frequency changes in genes involved in ion balance

Over the last 10,000 years, post-glacial stickleback populations have adapted to the different osmotic conditions found in freshwater (Spence et al. 2012). In this study, we found that seasonal isolation from the ocean led to repeated changes in allele frequency in many genes linked to ion balance. For example, genes such as *Nalcl* and *Wnk4* show signs of potential selection as shared outliers across multiple estuaries. *Nalcn* is a salt-sensing gene that was recently found to be rapidly evolving in saline-alkaline lake-dwelling fish (Tong and Li 2020). Similarly, *Wnk4* codes for an intracellular chloride sensor (Chen et al. 2019) implicated in salinity-tolerance in stickleback (Wang et al. 2014). Additionally, we found evidence of parallel selection on the gene *Ccny*. A recent epigenomic study linked *Ccny* to salinity adaptation, showing that *Ccny* is differentially methylated in stickleback populations along a gradient of decreasing salinity (Heckwolf et al. 2020). We also found evidence of selection on genes from the *Mapk* family. These genes are differentially expressed in fish in response to many environmental stressors (Mateus et al. 2017), including osmotic stress (Tse et al. 2011; Tian et al. 2019; reviewed by Kültz 2012).

We also found evidence of selection on genes for calcium balance, for example, the calcium sensing gene calmodulin (*calm1b*, Chin and Means 2000). Another gene we found to be putatively under seasonal parallel selection is the calcitonin receptor (*Calcr*), part of a family of genes known to regulate calcium homeostasis (Naot et al. 2019). Indeed, a gene from the same family, the calcitonin gene-related peptide type 1 receptor (*Calcrl*), has been implicated in salinity tolerance or osmoregulatory adaptation in postglacial stickleback populations (Kusakabe et al. 2017). These findings suggest a potential role for selection on these loci. However, to avoid false conclusions based on selective assessment of particular genes we analyzed the overall genetic functions of candidate genes.

### Temporal changes in bar-built estuaries likely select for ionic homeostasis

We focused on the top ten most significant GO terms in each outlier set and found functions related to ion transmembrane transport among candidate genes. Our findings are consistent with previous studies on freshwater adaptation that have documented divergence in genes linked to ion channels (e.g. sodium/potassium channels) during parallel adaptation to freshwater (DeFaveri et al. 2011; Jones et al. 2012a; Jones et al. 2012b; Gibbons et al. 2016; Gibbons et al. 2017; Hasan et al. 2017; Rudman et al. 2019; Heckwolf et al. 2020). Freshwater adaptation has occurred independently in a wide range of taxa through selection on genes involved in osmoregulation. For instance, annelids (Horn et al. 2019), arthropods (Lee et al. 2011), and fish (Velotta et al. 2016), including sculpin (Dennenmoser et al. 2016) and stickleback (Jones et al. 2012b; Kusakabe et al. 2016; Hasan et al. 2017), have all shown genetic changes in ion channel genes following colonization of freshwater habitats. Our findings are also in line with a recent study (Tong and Li 2020) on adaptation of *Gymnocypris przewalskii* to a saline-alkaline lake, which showed that rapidly evolving genes, those with an elevated rate of non-synonymous substitutions, are overwhelmingly involved in ion transport. Our findings of gene functions related to transmembrane ion transport could also be in part driven by changes in temperature over the time period sampled (during the summer months). Increases in temperature may disrupt osmoregulation. A study on estuary fish from California showed that experimentally exposing fish to higher temperature results in differential expression of some of the same genetic functions we found overrepresented among our candidate genes, including ion channel activity and extracellular matrix structural constituent (Jeffries et al. 2016). Thus, changes in temperature could amplify the osmoregulatory challenges experienced by populations responding to changes in salinity.

Freshwater adaptation in stickleback also could involve genetic changes to maintain calcium homeostasis (Gibbons et al. 2016). Calcium binding proteins play an important physiological role in maintaining calcium balance in fish (Evans et al. 2005). Calcium must be continuously absorbed from freshwater, which is hypoosmotic relative to fish plasma (Liem et al. 2001; reviewed by Evans et al. 2005). Fittingly, not only did we find consistent changes in allele frequency in genes relevant for calcium homeostasis, but we also found an overall enrichment of gene functions related to calcium ion binding. This provides additional evidence that genes for ion regulation are targets of selection during freshwater transitions that last from months to millennia.

## Conclusion

We found evidence of natural selection for osmoregulatory adaptation, likely brought into operation by seasonal changes in the ionic environment within estuaries. Repeated changes in allele frequency across estuaries suggests parallel selection is occurring, and the molecular functions targeted by selection have also been identified in retrospective studies of divergence between long-established marine and freshwater populations, highlighting the utility of this approach for making inferences about historical instances of adaptation. Our results are consistent with the idea that cyclical isolation and exposure to the ocean results in seasonally oscillating selection, although time-series data over multiple instances of sandbar formation is needed for confirmation. The threespine stickleback found in bar-built estuaries along coastal California thus provide the rare opportunity to study parallel selection in real-time, *in natura*.

## Acknowledgments

APH and RDHB were supported by NSERC Discovery Grants and Canada Research Chairs. Partial support for EPP was provided by the NOAA Cooperative Institute for Marine Ecosystems and Climate.

## Data accessibility

The data that support the findings of this manuscript will be uploaded to a publically accessible repository such as Sequence Read Archive (SRA) or Dryad. The custom bash scripts are available at https://github.com/garfring/scripts.

## Contributions

AP, EPP, APH, and RDHB conceived the study. AGE, AP, APH, and RDHB designed the methodological approach. BAW collected samples. AP performed the molecular work. AGE and AP performed the bioinformatics. AGE analyzed the data and created figures with assistance from TJT. AGE wrote the manuscript with input from all authors.

