## Supplemental Figures for "Using seasonal genomic changes to understand historical adaptation: parallel selection on stickleback in highly-variable estuaries"

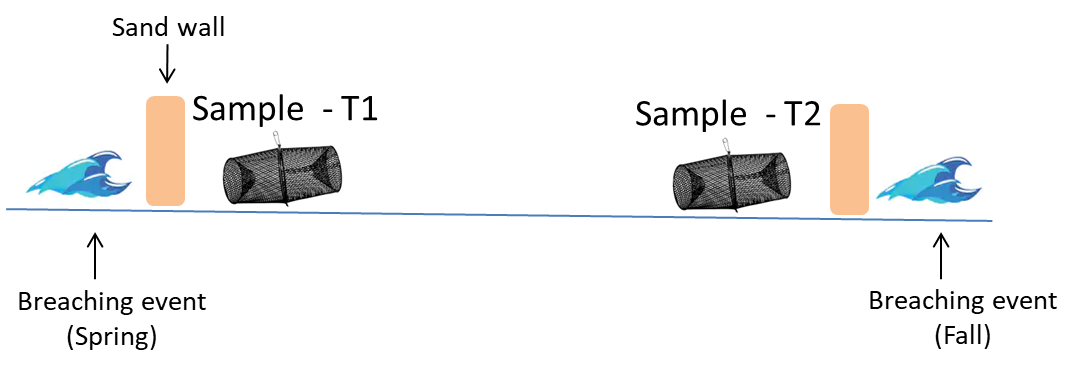
 Figure S1. Sampling of stickleback in each estuary occurred in between two breaching events. The first sample was taken shortly after the sandbar was rebuilt from a breaching event in the spring (T1). The second sampling time point was in the fall before the sandbar was broken in a new breaching event (T2). During this time while the sandbar is intact (T1 – T2), lentic estuaries transition from brackish to freshwater.

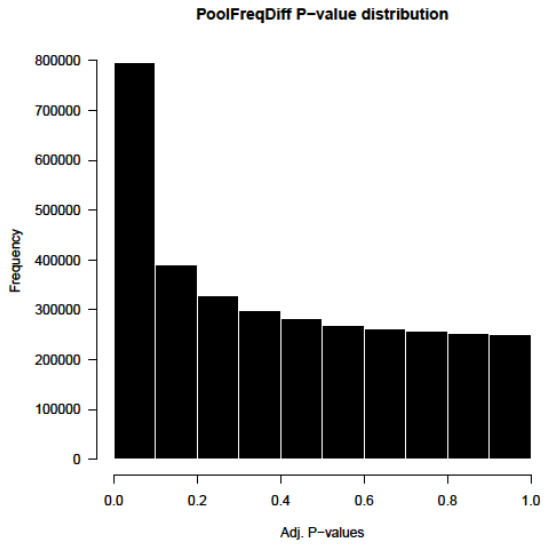

Figure S2. Histogram of P-values from the qGLM test (*PoolFreqDiff*) after adjusting for population structure using lambda (λ = 0.3) and prior to controlling for multiple hypothesis testing with *qvalue* program.

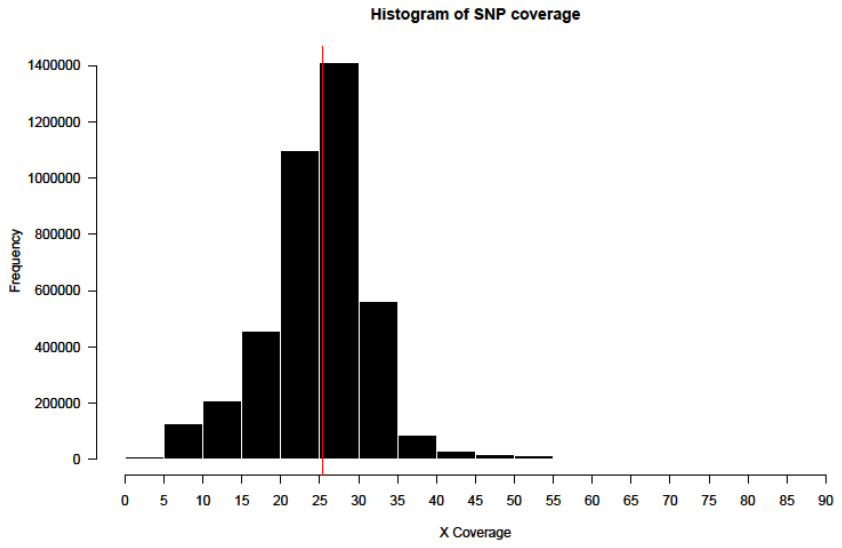
Figure S3. Histogram of minimum SNP coverage across estuaries. The average minimum coverage of 25.32 (SD = 6.96, range: 5 – 84) is denoted by the vertical red line.
