## Supplemental Table 1 for "Using seasonal genomic changes to understand historical adaptation: parallel selection on stickleback in highly-variable estuaries"

| Table S1. Site coordinates and size characteristics of each estuary. The size of each estuary was evaluated using Channel Area (CA), Permanently Flooded Area (CA + lakes, ponds, and side channels), and Total Wetland Area (maximum area that becomes seasonally flooded). | | | | | |
| --- | --- | --- | --- | --- | --- |
| Site | Latitude | Longitude | Channel Area (km^2^) | Permanently Flooded Area (km^2^) | Total Wetland Area (km^2^) |
| Scott Creek | 37.04 | -122.23 | 0.00635 | 0.01283 | 0.12509 |
| Laguna Creek | 36.984 | -122.154 | 0.02032 | 0.02185 | 0.05354 |
| Old Dairy Creek | 36.955 | -122.091 | 0.00004 | 4.00E-05 | 0.01331 |
| Younger Lagoon | 36.951 | -122.067 | 0.01081 | 0.01081 | 0.02092 |
| Lombardi Creek | 36.962 | -122.113 | 0.00117 | 0.00117 | 0.028 |
| Waddell Creek | 37.096 | -122.278 | 0.02004 | 0.0204 | 0.16908 |
