## Supplemental Table 2 for "Using seasonal genomic changes to understand historical adaptation: parallel selection on stickleback in highly-variable estuaries"

| Table S2. Permutation test (10,000 iterations) for an effect of estuary size on extent of parallelism. See Figure 1 for overall numbers n estuaries. | | | |
| --- | --- | --- | --- |
| SNPs outliers in *n* estuaries | Null expectation (95% quantile) | Observed number of SNPs | Empirical P-value |
| 3 (all small) | 1020 | 987 | 0.2723053 |
| 3 (all large) | 1020 | 966 | 0.5415162 |
