## Supplemental Table 3 for "Using seasonal genomic changes to understand historical adaptation: parallel selection on stickleback in highly-variable estuaries"

| Table S3. Genic F_ST_ outliers. | | |
| --- | --- | --- |
| Chromosome | Position | Overlapping gene |
| groupI | 85261 | col4a4 |
| groupI | 762830 | nf1a |
| groupI | 924144 | ubash3bb |
| groupI | 1210147 | sgsm2 |
| groupI | 1222323 | sgsm2 |
| groupI | 1356415 | myo1cb |
| groupI | 1356419 | myo1cb |
| groupI | 1612316 | si:ch211-276c2.2 |
| groupI | 2925222 | si:ch73-265h17.5 |
| groupI | 4497884 | slc33a1 |
| groupI | 5872569 | hip1 |
| groupI | 6494330 | DYNC2H1 |
| groupI | 6649525 | gria4a |
| groupI | 6931938 | FCHSD2 |
| groupI | 7877774 | limk1a |
| groupI | 9207104 | npas1 |
| groupI | 10040313 | ppm1lb |
| groupI | 10075425 | kpna4 |
| groupI | 10099523 | smc4 |
| groupI | 10176672 | schip1 |
| groupI | 12048484 | si:dkey-243i1.1 |
| groupI | 13702383 | frem2b |
| groupI | 14725652 | gab2 |
| groupI | 16276019 | NOVA1 |
| groupI | 16712599 | HNRNPUL1 |
| groupI | 17311060 | capn5a |
| groupI | 17977964 | zmp:0000001073 |
| groupI | 18039618 | zgc:101731 |
| groupI | 18383974 | aldh3a2a |
| groupI | 18418633 | ulk2 |
| groupI | 18462682 | usp32 |
| groupI | 18534255 | brip1 |
| groupI | 19617593 | ksr1b |
| groupI | 20253216 | kcnh7 |
| groupI | 21680480 | obsl1a |
| groupI | 22638012 | erbb4a |
| groupI | 25231590 | rap2ab |
| groupI | 26530632 | cd99 |
| groupI | 26558938 | dhrsx |
| groupI | 27013742 | hsf2bp |
| groupI | 27459733 | xdh |
| groupI | 27783813 | DES |
| groupII | 168168 | arnt2 |
| groupII | 484327 | ano9b |
| groupII | 1932057 | dpy19l3 |
| groupII | 1996614 | si:ch211-186j3.6 |
| groupII | 2023253 | si:ch211-186j3.6 |
| groupII | 2036781 | si:ch211-186j3.6 |
| groupII | 2687554 | CDH8 |
| groupII | 2716104 | CDH8 |
| groupII | 3450974 | ano1 |
| groupII | 3561457 | pcsk6 |
| groupII | 3632139 | pop4 |
| groupII | 5064721 | kif13ba |
| groupII | 6008546 | scml2 |
| groupII | 6310189 | itgb6 |
| groupII | 9489069 | plekha7b |
| groupII | 10829463 | zfhx3 |
| groupII | 11800158 | furina |
| groupII | 13939981 | kif26ba |
| groupII | 17868436 | nod2 |
| groupII | 17887414 | nkd1 |
| groupII | 18376096 | FBN1 |
| groupII | 18579021 | smad6a |
| groupII | 18962456 | KIF23 |
| groupII | 19168167 | CHRNA7 |
| groupII | 19168186 | CHRNA7 |
| groupII | 19357502 | MYO9A |
| groupII | 19511916 | neo1b |
| groupII | 19529077 | neo1b |
| groupII | 19723509 | mpped2a |
| groupII | 20020936 | prmt3 |
| groupII | 20154359 | NELL1 |
| groupII | 20163214 | NELL1 |
| groupII | 20164745 | NELL1 |
| groupII | 20223139 | NELL1 |
| groupII | 20859948 | kcnq1 |
| groupII | 21725408 | MEGF11 |
| groupII | 21898229 | otog |
| groupII | 22080909 | serinc4 |
| groupII | 22086970 | serinc4 |
| groupII | 22294374 | rbpms2b |
| groupII | 22538893 | fhod1 |
| groupII | 22632723 | wdr59 |
| groupIII | 995837 | ZNF521 |
| groupIII | 3255287 | rem2 |
| groupIII | 3973089 | zswim5 |
| groupIII | 4120217 | ctnnd2b |
| groupIII | 4593029 | rnf220a |
| groupIII | 6104099 | chd7 |
| groupIII | 6179834 | ca8 |
| groupIII | 7994872 | itga6b |
| groupIII | 8551306 | kcnn1a |
| groupIII | 8551317 | kcnn1a |
| groupIII | 11476793 | acvr2bb |
| groupIII | 12656591 | si:ch211-106h4.9 |
| groupIII | 13585006 | mib1 |
| groupIII | 13591612 | mib1 |
| groupIII | 15214945 | lig1 |
| groupIII | 15354432 | kirrel1b |
| groupIII | 16065943 | wrnip1 |
| groupIII | 16098153 | prkacbb |
| groupIII | 16139254 | si:ch1073-186i23.1 |
| groupIII | 16436611 | snai2 |
| groupIII | 16611642 | AGAP3 |
| groupIII | 16737274 | nol7 |
| groupIV | 177633 | rest |
| groupIV | 196289 | matr3l1.1 |
| groupIV | 1433258 | cxcl14 |
| groupIV | 2135893 | si:dkey-237h12.3 |
| groupIV | 2628302 | FSTL5 |
| groupIV | 2670202 | FSTL5 |
| groupIV | 2675175 | FSTL5 |
| groupIV | 2952251 | LIMCH1 |
| groupIV | 3199408 | fgf24 |
| groupIV | 3337184 | nudt6 |
| groupIV | 3337197 | nudt6 |
| groupIV | 3381695 | SPATA5 |
| groupIV | 5213913 | fermt3b |
| groupIV | 5490046 | efemp2a |
| groupIV | 6284022 | drp2 |
| groupIV | 8329956 | SLC16A2 |
| groupIV | 8424582 | mid2 |
| groupIV | 8424584 | mid2 |
| groupIV | 8436677 | mid2 |
| groupIV | 10492440 | spock1 |
| groupIV | 10961873 | ndst1a |
| groupIV | 11372731 | clk4a |
| groupIV | 12108815 | irg1l |
| groupIV | 12158965 | irg1l |
| groupIV | 13406322 | dctn4 |
| groupIV | 13406323 | dctn4 |
| groupIV | 13571693 | ctnna1 |
| groupIV | 13578848 | ctnna1 |
| groupIV | 14223518 | ablim3 |
| groupIV | 14225308 | ablim3 |
| groupIV | 14227500 | ablim3 |
| groupIV | 15005199 | TENM2 |
| groupIV | 15141467 | EFNB1 |
| groupIV | 15867851 | cnot7 |
| groupIV | 16007799 | gpc3 |
| groupIV | 16013845 | gpc3 |
| groupIV | 16104413 | brd8 |
| groupIV | 16358824 | si:ch73-49o8.1 |
| groupIV | 17630508 | irf2 |
| groupIV | 17942415 | ano6 |
| groupIV | 18234006 | grm8a |
| groupIV | 20995782 | SLC2A13 |
| groupIV | 21642308 | lrig3 |
| groupIV | 24122473 | scube1 |
| groupIV | 24182241 | scube1 |
| groupIV | 25003349 | OSBPL8 |
| groupIV | 25003475 | OSBPL8 |
| groupIV | 25068359 | zdhhc17 |
| groupIV | 25713675 | nxpe3 |
| groupIV | 25713799 | nxpe3 |
| groupIV | 25713900 | nxpe3 |
| groupIV | 25713920 | nxpe3 |
| groupIV | 25713960 | nxpe3 |
| groupIV | 25714242 | nxpe3 |
| groupIV | 25828152 | slc38a2 |
| groupIV | 25906741 | tm7sf3 |
| groupIV | 26006387 | setd7 |
| groupIV | 30100102 | hgfa |
| groupIV | 30228236 | snd1 |
| groupIV | 30776503 | anks1b |
| groupIV | 31092938 | si:ch73-352p4.8 |
| groupIV | 31109232 | TULP3 |
| groupIV | 31662074 | ppfia2 |
| groupIV | 31679576 | ppfia2 |
| groupIV | 31741054 | lin7a |
| groupIV | 31934666 | syt1a |
| groupIV | 32127569 | nup37 |
| groupIX | 610299 | rbfox2 |
| groupIX | 1667470 | klhl5 |
| groupIX | 1816241 | slit2 |
| groupIX | 2286243 | SLC24A3 |
| groupIX | 2377058 | sh3pxd2aa |
| groupIX | 4859265 | cyp3a65 |
| groupIX | 4859265 | cyp3a65 |
| groupIX | 4861626 | cyp3a65 |
| groupIX | 4861626 | cyp3a65 |
| groupIX | 4861730 | cyp3a65 |
| groupIX | 4861730 | cyp3a65 |
| groupIX | 4867044 | cyp3a65 |
| groupIX | 4867044 | cyp3a65 |
| groupIX | 4934834 | sdk1a |
| groupIX | 5208341 | si:dkeyp-9d4.3 |
| groupIX | 5959655 | tll1 |
| groupIX | 6407015 | lrba |
| groupIX | 7260230 | inpp4b |
| groupIX | 7263639 | inpp4b |
| groupIX | 7283138 | inpp4b |
| groupIX | 13978810 | smc3 |
| groupIX | 14967746 | pkd1a |
| groupIX | 15314114 | afmid |
| groupIX | 15728645 | xylt1 |
| groupIX | 18118315 | micall2b |
| groupIX | 18255428 | mala |
| groupIX | 18757824 | ppp3r1b |
| groupIX | 18940021 | tbce |
| groupIX | 18947442 | nid1b |
| groupIX | 19952391 | psen2 |
| groupV | 257699 | ADK |
| groupV | 262824 | ADK |
| groupV | 1246806 | lhpp |
| groupV | 1564568 | dock1 |
| groupV | 1990980 | opn4b |
| groupV | 3071211 | sdk2b |
| groupV | 3642949 | WNK4 |
| groupV | 4393370 | MYO1D |
| groupV | 4771292 | acsf2 |
| groupV | 4771292 | chad |
| groupV | 6028735 | SPAG9 |
| groupV | 6039736 | SPAG9 |
| groupV | 8333154 | cpn1 |
| groupV | 8362617 | dnmbp |
| groupV | 8371507 | dnmbp |
| groupV | 8526046 | ATRNL1 |
| groupV | 8788338 | cep55l |
| groupV | 8964836 | pcdh15b |
| groupV | 9676174 | aldoaa |
| groupV | 10260074 | ntn2 |
| groupV | 10355684 | im:7160594 |
| groupV | 10355685 | im:7160594 |
| groupV | 10472878 | ift140 |
| groupV | 11236388 | trrap |
| groupV | 11757092 | lipf |
| groupV | 12105748 | arg1 |
| groupVI | 65315 | pdzd8 |
| groupVI | 65320 | pdzd8 |
| groupVI | 535902 | piezo1 |
| groupVI | 628846 | degs1 |
| groupVI | 1469002 | STK32C |
| groupVI | 2224653 | btaf1 |
| groupVI | 2832952 | sorbs1 |
| groupVI | 2977696 | MYPN |
| groupVI | 3054537 | ATE1 |
| groupVI | 3315380 | cdh23 |
| groupVI | 3532982 | mlh1 |
| groupVI | 5214539 | jag1b |
| groupVI | 5561692 | macrod2 |
| groupVI | 6171741 | MCU |
| groupVI | 7263134 | adgrb3 |
| groupVI | 10695465 | pcdh15a |
| groupVI | 10720947 | pcdh15a |
| groupVI | 10720979 | pcdh15a |
| groupVI | 11196458 | add3b |
| groupVI | 11196522 | add3b |
| groupVI | 11972866 | dachc |
| groupVI | 15176816 | prkceb |
| groupVI | 15222130 | epas1b |
| groupVI | 16489290 | ddx43 |
| groupVI | 16698814 | entpd6 |
| groupVI | 16786362 | ppp2r5d |
| groupVII | 1533 | cnpy3 |
| groupVII | 734163 | hdac8 |
| groupVII | 1166395 | exoc6b |
| groupVII | 1527191 | supt16h |
| groupVII | 2069023 | arhgef11 |
| groupVII | 2262912 | zgc:55262 |
| groupVII | 2281161 | pacs1a |
| groupVII | 2927070 | efs |
| groupVII | 3225940 | tbc1d19 |
| groupVII | 3475396 | pcdh7b |
| groupVII | 3506539 | pcdh7b |
| groupVII | 4111347 | dvl2 |
| groupVII | 7590548 | slc8a4b |
| groupVII | 10022174 | NRXN2 |
| groupVII | 12416367 | wscd1b |
| groupVII | 12839808 | arhgap32b |
| groupVII | 14033304 | med13a |
| groupVII | 15251866 | akap10 |
| groupVII | 15655377 | gabra3 |
| groupVII | 17132205 | ar |
| groupVII | 17751863 | rb1 |
| groupVII | 18060796 | dpf2 |
| groupVII | 18428275 | grk1b |
| groupVII | 19030918 | asl |
| groupVII | 19037675 | asl |
| groupVII | 19041464 | asl |
| groupVII | 20697034 | map3k7cl |
| groupVII | 21450620 | slc47a1 |
| groupVII | 21450624 | slc47a1 |
| groupVII | 21464005 | slc47a1 |
| groupVII | 23608574 | gabrg2 |
| groupVII | 24285841 | zbtb16a |
| groupVII | 26201343 | gemin5 |
| groupVII | 26830639 | doc2b |
| groupVII | 27786221 | chm |
| groupVIII | 705499 | eps15l1a |
| groupVIII | 1366702 | slc1a8a |
| groupVIII | 1420692 | podn |
| groupVIII | 1439673 | slc1a7b |
| groupVIII | 1571263 | si:ch211-247n2.1 |
| groupVIII | 2019099 | lpp |
| groupVIII | 2070270 | lpp |
| groupVIII | 2140385 | aldh9a1a.1 |
| groupVIII | 3901055 | prrx1b |
| groupVIII | 5145150 | rgs8 |
| groupVIII | 7185822 | mpl |
| groupVIII | 7185827 | mpl |
| groupVIII | 9584441 | arid3a |
| groupVIII | 13574967 | acbd6 |
| groupVIII | 13585724 | acbd6 |
| groupVIII | 14756664 | shdb |
| groupVIII | 14764400 | shdb |
| groupVIII | 14966625 | trabd2b |
| groupVIII | 15943909 | ell |
| groupVIII | 16221404 | mbd3b |
| groupVIII | 16677624 | cbarpb |
| groupVIII | 16677625 | cbarpb |
| groupVIII | 17043307 | NCAN |
| groupVIII | 17173890 | hmha1b |
| groupVIII | 17173893 | hmha1b |
| groupVIII | 17448751 | dph5 |
| groupVIII | 17704983 | atp1b1a |
| groupVIII | 17715065 | nme7 |
| groupVIII | 17953010 | cyfip1 |
| groupVIII | 17959965 | cyfip1 |
| groupVIII | 17966434 | cyfip1 |
| groupVIII | 18218521 | AP1S2 |
| groupVIII | 18228657 | SHROOM2 |
| groupVIII | 18311767 | nek1 |
| groupVIII | 18536876 | scfd2 |
| groupVIII | 19036986 | uap1 |
| groupVIII | 19206380 | echdc2 |
| groupVIII | 19206380 | zyg11 |
| groupX | 1031088 | pou2f2a |
| groupX | 1434269 | rnf19b |
| groupX | 2022424 | xkr8.2 |
| groupX | 2236081 | atp9b |
| groupX | 2372157 | pleca |
| groupX | 3397057 | med30 |
| groupX | 3946811 | TGFBR2 |
| groupX | 6434188 | AKAP9 |
| groupX | 6467748 | krit1 |
| groupX | 6959237 | efna1a |
| groupX | 8346636 | elmo1 |
| groupX | 8912520 | calcr |
| groupX | 9268603 | col1a2 |
| groupX | 9313919 | ppp1r9a |
| groupX | 9559058 | igf2bp3 |
| groupX | 10643412 | col16a1 |
| groupX | 10651654 | col16a1 |
| groupX | 11313196 | gstr |
| groupX | 11411476 | arid1ab |
| groupX | 12140845 | MANEAL |
| groupX | 12376549 | top2b |
| groupX | 13158201 | ST3GAL1 |
| groupX | 13158217 | ST3GAL1 |
| groupX | 13346262 | KHDRBS3 |
| groupX | 15057778 | vps52 |
| groupXI | 2784937 | zgc:63569 |
| groupXI | 3037606 | nsfa |
| groupXI | 3181135 | cpt1cb |
| groupXI | 4127932 | si:ch1073-322p19.1 |
| groupXI | 5242096 | zgc:113411 |
| groupXI | 6319058 | vat1 |
| groupXI | 8141369 | dnm2a |
| groupXI | 9138672 | cant1a |
| groupXI | 9863208 | coro7 |
| groupXI | 9892926 | coro7 |
| groupXI | 12085729 | gtpbp1 |
| groupXI | 12097682 | sgsm3 |
| groupXI | 12967327 | sec14l1 |
| groupXI | 14638719 | LLGL1 |
| groupXI | 14692631 | drg2 |
| groupXI | 14862190 | cacng3b |
| groupXI | 14868809 | cacng3b |
| groupXI | 15669080 | CACNG2 |
| groupXI | 15820278 | med15 |
| groupXI | 15853154 | btr30 |
| groupXI | 16239896 | mfsd11 |
| groupXII | 173234 | prpf6 |
| groupXII | 219262 | nkain4 |
| groupXII | 778543 | magi1a |
| groupXII | 1064128 | nr2c2 |
| groupXII | 1390432 | adcy6a |
| groupXII | 1582014 | pfkfb2a |
| groupXII | 1607297 | scube3 |
| groupXII | 1661807 | ANKS1A |
| groupXII | 1665691 | ANKS1A |
| groupXII | 2054151 | mitfb |
| groupXII | 3965736 | ntsr1 |
| groupXII | 4021290 | slco4a1 |
| groupXII | 4137490 | ralgapb |
| groupXII | 4167543 | prex1 |
| groupXII | 4167960 | prex1 |
| groupXII | 4822717 | plxna1a |
| groupXII | 5159212 | agrn |
| groupXII | 6460320 | dnajc16 |
| groupXII | 6796067 | sema3h |
| groupXII | 9692047 | noc2l |
| groupXII | 9836616 | rerea |
| groupXII | 13491489 | hsd17b10 |
| groupXII | 13764043 | cs |
| groupXII | 14138697 | plxna2 |
| groupXII | 14143778 | plxna2 |
| groupXII | 14989561 | acap3a |
| groupXII | 15764340 | ythdf1 |
| groupXII | 16832979 | ephb2a |
| groupXII | 16833056 | ephb2a |
| groupXII | 17501954 | skia |
| groupXII | 17502365 | skia |
| groupXII | 17531731 | skia |
| groupXII | 18301421 | ccdc22 |
| groupXIII | 1000849 | zgc:64051 |
| groupXIII | 1048433 | gtf2h2 |
| groupXIII | 1951818 | ndufs4 |
| groupXIII | 2619063 | rad17 |
| groupXIII | 3217071 | NSMF |
| groupXIII | 4836805 | DMXL1 |
| groupXIII | 4842212 | DMXL1 |
| groupXIII | 5081419 | loxhd1b |
| groupXIII | 6211357 | zswim6 |
| groupXIII | 6532396 | smarcad1a |
| groupXIII | 6532401 | smarcad1a |
| groupXIII | 6629991 | grid2 |
| groupXIII | 7469484 | ndc80 |
| groupXIII | 9245655 | p2rx2 |
| groupXIII | 9245657 | p2rx2 |
| groupXIII | 11053039 | fam172a |
| groupXIII | 11293846 | mctp1a |
| groupXIII | 12344504 | zgc:77112 |
| groupXIII | 13798927 | angptl2b |
| groupXIII | 14334245 | ncor2 |
| groupXIII | 14747990 | MSI1 |
| groupXIII | 14892645 | htr7c |
| groupXIII | 15336090 | chfr |
| groupXIII | 16056038 | hk2 |
| groupXIII | 16098803 | mxd1 |
| groupXIII | 16372407 | si:dkey-91m11.5 |
| groupXIII | 16374537 | si:dkey-91m11.5 |
| groupXIII | 16554401 | citb |
| groupXIII | 17713078 | col27a1b |
| groupXIII | 18861948 | cds1 |
| groupXIII | 19081696 | adamts3 |
| groupXIII | 19113887 | adamts3 |
| groupXIII | 19258058 | dmrt1 |
| groupXIII | 19368639 | atp8b5a |
| groupXIII | 19389293 | cnnm4b |
| groupXIV | 191297 | apc |
| groupXIV | 670391 | adamts6 |
| groupXIV | 902522 | pde8b |
| groupXIV | 1233560 | pik3r1 |
| groupXIV | 1676000 | si:dkey-226m8.10 |
| groupXIV | 2022655 | col5a1 |
| groupXIV | 3450849 | plpp1a |
| groupXIV | 6336247 | WDR70 |
| groupXIV | 7479984 | rc3h2 |
| groupXIV | 7560515 | strbp |
| groupXIV | 7691338 | cercam |
| groupXIV | 7691412 | cercam |
| groupXIV | 7724003 | dbh |
| groupXIV | 7724067 | dbh |
| groupXIV | 7724068 | dbh |
| groupXIV | 7928407 | cdc37l1 |
| groupXIV | 8230710 | si:ch73-287m6.1 |
| groupXIV | 8317774 | si:dkey-112m2.1 |
| groupXIV | 8627095 | sfswap |
| groupXIV | 8802409 | mmp17a |
| groupXIV | 9438357 | GRIN3A |
| groupXIV | 10128595 | cntfr |
| groupXIV | 11047129 | man1b1a |
| groupXIV | 11073386 | whrna |
| groupXIV | 11394032 | tsc1b |
| groupXIV | 11394034 | tsc1b |
| groupXIV | 13226808 | WDR7 |
| groupXIV | 13256683 | ccng2 |
| groupXIV | 13558515 | dcc |
| groupXIV | 13562065 | dcc |
| groupXIV | 13786838 | fbxw2 |
| groupXIV | 14341827 | slc25a25b |
| groupXIV | 14530148 | fpgs |
| groupXIV | 14530148 | cdk9 |
| groupXIV | 14656013 | crata |
| groupXIX | 155399 | AGBL1 |
| groupXIX | 535966 | zgc:158366 |
| groupXIX | 696909 | prmt7 |
| groupXIX | 696910 | prmt7 |
| groupXIX | 767590 | smpd3 |
| groupXIX | 855929 | gins3 |
| groupXIX | 1177562 | btbd10a |
| groupXIX | 1497498 | ppp6r3 |
| groupXIX | 1604388 | ces3 |
| groupXIX | 1605843 | ces3 |
| groupXIX | 2101576 | siah1 |
| groupXIX | 2111122 | siah1 |
| groupXIX | 2123707 | lonp2 |
| groupXIX | 2486385 | myh7bb |
| groupXIX | 2693940 | necab2 |
| groupXIX | 2768433 | mbtps1 |
| groupXIX | 2910599 | ranbp10 |
| groupXIX | 2919345 | ranbp10 |
| groupXIX | 3058100 | klhdc4 |
| groupXIX | 3165547 | fa2h |
| groupXIX | 3696916 | tead1a |
| groupXIX | 3729185 | parvaa |
| groupXIX | 3729194 | parvaa |
| groupXIX | 3729222 | parvaa |
| groupXIX | 4601439 | mon2 |
| groupXIX | 4603471 | mon2 |
| groupXIX | 6187781 | dhtkd1 |
| groupXIX | 6187938 | dhtkd1 |
| groupXIX | 6936668 | LRRC10B |
| groupXIX | 6956001 | syt7a |
| groupXIX | 6956003 | syt7a |
| groupXIX | 6956013 | syt7a |
| groupXIX | 7341537 | ano3 |
| groupXIX | 7375690 | clpxb |
| groupXIX | 7375691 | clpxb |
| groupXIX | 7375693 | clpxb |
| groupXIX | 7375707 | clpxb |
| groupXIX | 7432798 | MYO5A |
| groupXIX | 7582137 | tjp1b |
| groupXIX | 8714499 | lamb1a |
| groupXIX | 8763053 | frs2b |
| groupXIX | 8763087 | frs2b |
| groupXIX | 8860525 | gpia |
| groupXIX | 9325487 | tnni2a.3 |
| groupXIX | 9338343 | lsp1 |
| groupXIX | 9356436 | tnnt3a |
| groupXIX | 10225695 | asz1 |
| groupXIX | 10237889 | asz1 |
| groupXIX | 10844748 | alx1 |
| groupXIX | 11450107 | RELN |
| groupXIX | 11467173 | RELN |
| groupXIX | 11656169 | PDE3A |
| groupXIX | 12158600 | tln2b |
| groupXIX | 12170304 | tln2b |
| groupXIX | 12396182 | roraa |
| groupXIX | 13231403 | osbpl5 |
| groupXIX | 13355991 | mrpl23 |
| groupXIX | 13420421 | cpt1ab |
| groupXIX | 13760083 | ldha |
| groupXIX | 13779758 | hrasb |
| groupXIX | 13783376 | hrasb |
| groupXIX | 13783774 | hrasb |
| groupXIX | 13916379 | LDLRAD3 |
| groupXIX | 13916381 | LDLRAD3 |
| groupXIX | 13916396 | LDLRAD3 |
| groupXIX | 13916400 | LDLRAD3 |
| groupXIX | 14035773 | pamr1 |
| groupXIX | 14035877 | pamr1 |
| groupXIX | 14037793 | pamr1 |
| groupXIX | 14037805 | pamr1 |
| groupXIX | 14679371 | banp |
| groupXIX | 14893013 | rassf8a |
| groupXIX | 14934058 | itpr2 |
| groupXIX | 15004158 | nr1h4 |
| groupXIX | 15235216 | magi2b |
| groupXIX | 15260020 | magi2b |
| groupXIX | 15348919 | prr5a |
| groupXIX | 15348921 | prr5a |
| groupXIX | 16083573 | rbm28 |
| groupXIX | 16675756 | lmf2a |
| groupXIX | 16917923 | IGF1R |
| groupXIX | 16917928 | IGF1R |
| groupXIX | 18144486 | hipk2 |
| groupXIX | 18154488 | hipk2 |
| groupXIX | 18555879 | tmtc2b |
| groupXIX | 18566263 | tmtc2b |
| groupXIX | 18566268 | tmtc2b |
| groupXIX | 18669924 | bicd1a |
| groupXIX | 18676323 | bicd1a |
| groupXIX | 18808896 | ada2a |
| groupXIX | 18979705 | zgc:173742 |
| groupXIX | 18981760 | zgc:173742 |
| groupXIX | 19002277 | mlycd |
| groupXIX | 19063381 | cdh13 |
| groupXIX | 19529735 | e2f4 |
| groupXIX | 19532706 | e2f4 |
| groupXIX | 19799949 | cbfb |
| groupXIX | 19799953 | cbfb |
| groupXIX | 19868929 | GNAO1 |
| groupXV | 420963 | slc4a11 |
| groupXV | 623475 | fgfr3 |
| groupXV | 878013 | mark3a |
| groupXV | 1006490 | BRF1 |
| groupXV | 1634088 | galnt16 |
| groupXV | 2227371 | RGS6 |
| groupXV | 2472773 | FMN1 |
| groupXV | 2715782 | akt1 |
| groupXV | 3071522 | ZDHHC14 |
| groupXV | 3660135 | SCAF8 |
| groupXV | 4347319 | ldlrap1b |
| groupXV | 4347322 | ldlrap1b |
| groupXV | 5993917 | ltk |
| groupXV | 6042640 | si:dkey-13p1.4 |
| groupXV | 6245246 | kidins220a |
| groupXV | 6929983 | znf292a |
| groupXV | 8210359 | crlf1a |
| groupXV | 10439279 | snap23.1 |
| groupXV | 10856058 | slc25a21 |
| groupXV | 11927596 | pacs2 |
| groupXV | 11964154 | actn1 |
| groupXV | 14239298 | babam2 |
| groupXV | 15455501 | setd3 |
| groupXV | 16025397 | kcnk5a |
| groupXVI | 3648993 | sestd1 |
| groupXVI | 4150750 | RAMP1 |
| groupXVI | 6455386 | slc40a1 |
| groupXVI | 6455392 | slc40a1 |
| groupXVI | 6455500 | slc40a1 |
| groupXVI | 6711719 | cmss1 |
| groupXVI | 7012607 | rcbtb1 |
| groupXVI | 7734394 | appb |
| groupXVI | 8201834 | GSK3B |
| groupXVI | 8543530 | nqo1 |
| groupXVI | 9578110 | ADAM23 |
| groupXVI | 11786779 | trpm2 |
| groupXVI | 12645970 | erbb4b |
| groupXVI | 14401010 | stxbp5l |
| groupXVI | 15927404 | lrch1 |
| groupXVI | 15927437 | lrch1 |
| groupXVI | 15994142 | sh3bp4 |
| groupXVI | 16098029 | caska |
| groupXVI | 16558988 | abcb6b |
| groupXVI | 17584426 | plcl1 |
| groupXVII | 1379422 | ATP6AP1L |
| groupXVII | 1677561 | atp2b2 |
| groupXVII | 2321752 | ephb2b |
| groupXVII | 3617667 | si:dkey-238f9.1 |
| groupXVII | 6303657 | znf362a |
| groupXVII | 6999690 | hdac7b |
| groupXVII | 10053721 | cdh4 |
| groupXVII | 10420063 | zgc:92107 |
| groupXVII | 10447506 | ndrg3a |
| groupXVII | 10987464 | phactr3b |
| groupXVII | 11679213 | cfap57 |
| groupXVII | 11917198 | fbln2 |
| groupXVII | 12211780 | CPNE9 |
| groupXVII | 12227031 | mtmr14 |
| groupXVII | 12620833 | cyb561d1 |
| groupXVII | 12620834 | cyb561d1 |
| groupXVII | 13731462 | pdzrn3b |
| groupXVIII | 724190 | cd2ap |
| groupXVIII | 747441 | tnfrsf21 |
| groupXVIII | 1298178 | cdc42bpab |
| groupXVIII | 2016697 | calm1b |
| groupXVIII | 2317408 | nrxn3b |
| groupXVIII | 2334678 | nrxn3b |
| groupXVIII | 2955853 | cgrrf1 |
| groupXVIII | 3355320 | nt5dc1 |
| groupXVIII | 4709916 | cdk19 |
| groupXVIII | 5385454 | tbc1d32 |
| groupXVIII | 9208673 | mrap2b |
| groupXVIII | 9398199 | matn3a |
| groupXVIII | 10169758 | fdft1 |
| groupXVIII | 10708381 | snx17 |
| groupXVIII | 10965226 | eya4 |
| groupXVIII | 10965334 | eya4 |
| groupXVIII | 11368094 | nbas |
| groupXVIII | 13449171 | si:ch1073-155h21.1 |
| groupXVIII | 13650635 | runx2b |
| groupXVIII | 13768203 | fkbp1b |
| groupXVIII | 14165771 | PLCB1 |
| groupXVIII | 15030531 | pigh |
| groupXVIII | 15431671 | rnf217 |
| groupXVIII | 16173901 | mthfd1a |
| groupXX | 83790 | dennd4b |
| groupXX | 174683 | pik3r4 |
| groupXX | 399080 | IQGAP3 |
| groupXX | 563010 | myo1eb |
| groupXX | 2009335 | ano10a |
| groupXX | 2978274 | RIMS2 |
| groupXX | 6277129 | cobl |
| groupXX | 6934056 | nrsn1 |
| groupXX | 7470424 | cdkal1 |
| groupXX | 7484682 | cdkal1 |
| groupXX | 7606880 | COLEC10 |
| groupXX | 7606890 | COLEC10 |
| groupXX | 7686849 | CARMIL1 |
| groupXX | 7686851 | CARMIL1 |
| groupXX | 7933731 | col14a1a |
| groupXX | 8793862 | si:ch73-380l3.2 |
| groupXX | 8814434 | si:ch73-380l3.2 |
| groupXX | 10265744 | steap4 |
| groupXX | 10518699 | tmem245 |
| groupXX | 10909957 | pafah1b3 |
| groupXX | 11117793 | cicb |
| groupXX | 11608973 | lpcat3 |
| groupXX | 11723404 | zgc:92912 |
| groupXX | 12088534 | foxj2 |
| groupXX | 12819596 | crabp2a |
| groupXX | 14703661 | VWDE |
| groupXX | 16834661 | MAN1C1 |
| groupXX | 17207593 | fhl3a |
| groupXX | 17772190 | smpdl3b |
| groupXX | 18664829 | ascc3 |
| groupXX | 18664871 | ascc3 |
| groupXX | 18939655 | cdk14 |
| groupXXI | 1068422 | PARD3 |
| groupXXI | 1323162 | ccny |
| groupXXI | 1861396 | adarb2 |
| groupXXI | 2183125 | abca4a |
| groupXXI | 2308001 | col11a1a |
| groupXXI | 4929720 | zfhx4 |
| groupXXI | 5928041 | pola1 |
| groupXXI | 5928082 | pola1 |
| groupXXI | 6849448 | eya1 |
| groupXXI | 9058787 | lmbr1 |
| groupXXI | 9259242 | esyt2b |
| groupXXI | 9485833 | COLEC12 |
| groupXXI | 9961350 | asap1b |
| groupXXI | 10093284 | puf60b |
| groupXXI | 10292686 | gli3 |
| groupXXI | 10294135 | gli3 |
| groupXXI | 11100657 | gpr158a |
