## Supplemental Table 4 for "Using seasonal genomic changes to understand historical adaptation: parallel selection on stickleback in highly-variable estuaries"

| Table S4. qGLM outliers (FDR = 0.01%) mapped to genes. | | |
| --- | --- | --- |
| Chromosome | Position | Overlapping gene |
| groupI | 194284 | ANO7 |
| groupI | 194297 | ANO7 |
| groupI | 1356417 | myo1cb |
| groupI | 2079573 | numbl |
| groupI | 2079695 | numbl |
| groupI | 2225424 | CEP164 |
| groupI | 2225425 | CEP164 |
| groupI | 2294609 | MPZL2 |
| groupI | 2921889 | si:ch73-265h17.5 |
| groupI | 4359566 | dscamb |
| groupI | 4486100 | RPS6KB1 |
| groupI | 6503833 | DYNC2H1 |
| groupI | 6652237 | gria4a |
| groupI | 8623856 | sik2b |
| groupI | 9151616 | slc1a5 |
| groupI | 11028941 | ROBO3 |
| groupI | 11997404 | pitpnm3 |
| groupI | 12205295 | synrg |
| groupI | 14268117 | cog6 |
| groupI | 16878384 | aplp2 |
| groupI | 17023775 | MAP4K1 |
| groupI | 18834669 | bcas3 |
| groupI | 19552437 | akap1b |
| groupI | 19761643 | PIPOX |
| groupI | 19761644 | PIPOX |
| groupI | 19777379 | PIPOX |
| groupI | 20058982 | ical1 |
| groupI | 20075909 | bmpr2a |
| groupI | 22857862 | kpnb3 |
| groupI | 24615905 | ARHGAP6 |
| groupI | 24647536 | ARHGAP6 |
| groupI | 26035523 | pudp |
| groupI | 26123756 | nlgn4 |
| groupI | 27198465 | gucy1b2 |
| groupI | 27795801 | map3k2 |
| groupI | 27892870 | ahr1b |
| groupII | 64906 | cemip |
| groupII | 72713 | cemip |
| groupII | 1386127 | znf536 |
| groupII | 1895302 | znf507 |
| groupII | 1976574 | si:ch211-186j3.6 |
| groupII | 2069386 | si:ch211-186j3.6 |
| groupII | 2482071 | adgrg1 |
| groupII | 3096000 | adamts18 |
| groupII | 3386926 | zgc:153993 |
| groupII | 3386930 | zgc:153993 |
| groupII | 3810111 | ccdc102a |
| groupII | 4307052 | CSK |
| groupII | 5507405 | itfg1 |
| groupII | 5507408 | itfg1 |
| groupII | 5646182 | wtip |
| groupII | 6072158 | ace2 |
| groupII | 7405938 | pcdh17 |
| groupII | 7845640 | GPC5 |
| groupII | 9205254 | sord |
| groupII | 9352682 | dagla |
| groupII | 9360871 | dagla |
| groupII | 9464226 | plekha7b |
| groupII | 9484845 | plekha7b |
| groupII | 9485992 | plekha7b |
| groupII | 11339059 | aldh1a2 |
| groupII | 11413867 | cgnl1 |
| groupII | 11486721 | znf280d |
| groupII | 11486724 | znf280d |
| groupII | 12887822 | elp4 |
| groupII | 13091572 | CSMD1 |
| groupII | 14027788 | fzd3b |
| groupII | 14759789 | mdga1 |
| groupII | 16360482 | ugt5f1 |
| groupII | 18426961 | SECISBP2L |
| groupII | 19317537 | gramd2aa |
| groupII | 19529077 | neo1b |
| groupII | 21905813 | otog |
| groupII | 22473755 | PLEKHG4 |
| groupII | 22516645 | fhod1 |
| groupII | 22640762 | wdr59 |
| groupIII | 109352 | col6a1 |
| groupIII | 532867 | PHACTR1 |
| groupIII | 1339603 | dpp6b |
| groupIII | 1349077 | arhgap21b |
| groupIII | 1357304 | arhgap21b |
| groupIII | 2025815 | eif2b3 |
| groupIII | 2028072 | eif2b3 |
| groupIII | 2325889 | gpc5c |
| groupIII | 3187533 | carmil3 |
| groupIII | 3245336 | rem2 |
| groupIII | 4944608 | crispld1b |
| groupIII | 7401013 | onecut3b |
| groupIII | 8295261 | upf1 |
| groupIII | 8875390 | rpap2 |
| groupIII | 8876148 | rpap2 |
| groupIII | 9580896 | MIER1 |
| groupIII | 10334379 | abi1b |
| groupIII | 11368126 | PTPRM |
| groupIII | 11395637 | PTPRM |
| groupIII | 11742815 | vav3b |
| groupIII | 12115738 | PTPRF |
| groupIII | 12115741 | PTPRF |
| groupIII | 12458632 | prkci |
| groupIII | 12527450 | samd7 |
| groupIII | 12735410 | cacnb2b |
| groupIII | 12915935 | lpin2 |
| groupIII | 13274093 | b4galt2 |
| groupIII | 13585312 | mib1 |
| groupIII | 13881945 | pth1ra |
| groupIII | 15164917 | kcnj9 |
| groupIII | 15202767 | lig1 |
| groupIII | 16710805 | cdk5 |
| groupIII | 16767818 | retreg1 |
| groupIII | 16767819 | retreg1 |
| groupIII | 16768113 | retreg1 |
| groupIV | 215897 | tgfbi |
| groupIV | 1019927 | gpc2 |
| groupIV | 2243320 | pdgfc |
| groupIV | 2601190 | FSTL5 |
| groupIV | 2620952 | FSTL5 |
| groupIV | 2725245 | ldb2a |
| groupIV | 3344338 | SPATA5 |
| groupIV | 3518208 | ANKRD50 |
| groupIV | 3897816 | PCGF3 |
| groupIV | 4522355 | glra1 |
| groupIV | 4950162 | mgat4b |
| groupIV | 4950163 | mgat4b |
| groupIV | 5185177 | fermt3b |
| groupIV | 5567545 | slc25a43 |
| groupIV | 5872589 | mcf2a |
| groupIV | 5875137 | mcf2a |
| groupIV | 6162198 | arhgef6 |
| groupIV | 7962155 | BEND4 |
| groupIV | 8529654 | si:ch211-220e11.3 |
| groupIV | 11356900 | RASGEF1C |
| groupIV | 11363916 | RASGEF1C |
| groupIV | 12568372 | nlgn3a |
| groupIV | 14910821 | TENM2 |
| groupIV | 14931022 | TENM2 |
| groupIV | 14931034 | TENM2 |
| groupIV | 15087182 | stag2b |
| groupIV | 15552599 | si:ch211-26b3.4 |
| groupIV | 16755575 | si:ch211-159i8.4 |
| groupIV | 18294419 | grm8a |
| groupIV | 22466149 | ptprz1b |
| groupIV | 22904945 | mkrn1 |
| groupIV | 22927334 | tmem178b |
| groupIV | 23122945 | tmtc1 |
| groupIV | 23122946 | tmtc1 |
| groupIV | 23837305 | hspa14 |
| groupIV | 24380863 | mkln1 |
| groupIV | 25592793 | ANO4 |
| groupIV | 28487084 | wnt5b |
| groupIV | 28514992 | wnt5b |
| groupIV | 28607445 | ERC1 |
| groupIV | 28698922 | wnk1b |
| groupIV | 29365747 | chchd3a |
| groupIV | 29383782 | chchd3a |
| groupIV | 29713181 | iqsec3a |
| groupIV | 30041546 | cacna2d1a |
| groupIV | 30161642 | snd1 |
| groupIV | 31019585 | atp6v1e1b |
| groupIV | 31163979 | ptn |
| groupIV | 32039274 | btbd11a |
| groupIV | 32124668 | parpbp |
| groupIV | 32379845 | acot16 |
| groupIX | 1233489 | pcdh7a |
| groupIX | 1519539 | pax5 |
| groupIX | 2371846 | sh3pxd2aa |
| groupIX | 4059070 | kdm2aa |
| groupIX | 5665301 | fat1a |
| groupIX | 5680724 | fat1a |
| groupIX | 5957375 | tll1 |
| groupIX | 6033487 | prom1b |
| groupIX | 6789945 | pou4f2 |
| groupIX | 7312774 | rnf150a |
| groupIX | 8002116 | lef1 |
| groupIX | 8395482 | mxd4 |
| groupIX | 9189845 | ctnna2 |
| groupIX | 10822865 | wbp1lb |
| groupIX | 11188437 | dok7 |
| groupIX | 12341680 | NAA15 |
| groupIX | 12861345 | prkca |
| groupIX | 12938477 | helz |
| groupIX | 15454806 | gcdha |
| groupIX | 15456561 | gcdha |
| groupIX | 15578032 | C16orf45 |
| groupIX | 15892809 | ccdc85al |
| groupIX | 17068902 | gga3 |
| groupIX | 18849813 | strn |
| groupIX | 19216545 | si:ch211-133l11.10 |
| groupIX | 19233714 | USP34 |
| groupIX | 19520170 | myom2a |
| groupIX | 19895755 | gpr108 |
| groupIX | 20141599 | zgc:109889 |
| groupV | 249160 | ADK |
| groupV | 1028846 | CDHR1 |
| groupV | 1657571 | ptprea |
| groupV | 1873318 | mta3 |
| groupV | 2549872 | prph2a |
| groupV | 3530315 | TANC2 |
| groupV | 3652480 | WNK4 |
| groupV | 3652485 | WNK4 |
| groupV | 3652488 | WNK4 |
| groupV | 3985769 | tmem106a |
| groupV | 3987676 | tmem106a |
| groupV | 4610493 | hoxb5b |
| groupV | 4610517 | hoxb5b |
| groupV | 4610526 | hoxb5b |
| groupV | 4610538 | hoxb5b |
| groupV | 4610541 | hoxb5b |
| groupV | 4610543 | hoxb5b |
| groupV | 4610559 | hoxb5b |
| groupV | 4610561 | hoxb5b |
| groupV | 4610562 | hoxb5b |
| groupV | 4610573 | hoxb5b |
| groupV | 4661278 | fbrs |
| groupV | 6480734 | cacna1g |
| groupV | 6783702 | SLC39A11 |
| groupV | 6829574 | usp22 |
| groupV | 8172811 | DNAJC7 |
| groupV | 8257791 | VAT1 |
| groupV | 8931495 | pcdh15b |
| groupV | 10375451 | MPRIP |
| groupV | 11239174 | trrap |
| groupVI | 360048 | cdh30 |
| groupVI | 362892 | cdh30 |
| groupVI | 383491 | cdh30 |
| groupVI | 674490 | polr1c |
| groupVI | 674493 | polr1c |
| groupVI | 2817311 | sorbs1 |
| groupVI | 2939368 | bag3 |
| groupVI | 2970516 | MYPN |
| groupVI | 3188922 | wdr11 |
| groupVI | 3776332 | fbxw4 |
| groupVI | 4182520 | b3gat2 |
| groupVI | 4343168 | PTK7 |
| groupVI | 4725102 | sipa1l2 |
| groupVI | 7082913 | fam135a |
| groupVI | 7088860 | fam135a |
| groupVI | 7088868 | fam135a |
| groupVI | 7534195 | khdrbs2 |
| groupVI | 7853672 | gclc |
| groupVI | 8201351 | WDFY4 |
| groupVI | 10059688 | kcnma1a |
| groupVI | 10125565 | kcnma1a |
| groupVI | 11042169 | neurl1ab |
| groupVI | 11042202 | neurl1ab |
| groupVI | 11042204 | neurl1ab |
| groupVI | 11573676 | adam12 |
| groupVI | 12204800 | bicc1a |
| groupVI | 12209702 | bicc1a |
| groupVI | 12212527 | bicc1a |
| groupVI | 13821175 | CCDC88A |
| groupVI | 13944537 | kcnh1a |
| groupVI | 13976992 | cyp3c3 |
| groupVI | 14016829 | USP54 |
| groupVI | 14121472 | kif20ba |
| groupVI | 14490864 | SPTBN1 |
| groupVI | 15543216 | nckap1 |
| groupVI | 15684936 | napba |
| groupVI | 15685231 | napba |
| groupVI | 15725087 | mcm8 |
| groupVI | 16729856 | ninl |
| groupVI | 16887743 | ubr2 |
| groupVII | 170884 | tec |
| groupVII | 1152920 | exoc6b |
| groupVII | 1633750 | si:dkey-46i9.6 |
| groupVII | 1633751 | si:dkey-46i9.6 |
| groupVII | 1853284 | ltb4r2a |
| groupVII | 2060777 | arhgef11 |
| groupVII | 2070514 | arhgef11 |
| groupVII | 2264587 | zgc:55262 |
| groupVII | 2491155 | intu |
| groupVII | 3115871 | chd3 |
| groupVII | 7279206 | pcxb |
| groupVII | 8915527 | fgf11a |
| groupVII | 9372934 | ndufs2 |
| groupVII | 10491101 | cblb |
| groupVII | 10827138 | picalma |
| groupVII | 10870723 | dlg2 |
| groupVII | 10939050 | dlg2 |
| groupVII | 10985051 | dlg2 |
| groupVII | 10987012 | dlg2 |
| groupVII | 10995435 | dlg2 |
| groupVII | 10995450 | dlg2 |
| groupVII | 11022315 | dlg2 |
| groupVII | 11496027 | ntm |
| groupVII | 13128851 | dscama |
| groupVII | 15498647 | slc8a4a |
| groupVII | 16924111 | fgf11b |
| groupVII | 17068916 | TBC1D8B |
| groupVII | 18010043 | slc25a35 |
| groupVII | 19037705 | asl |
| groupVII | 19046228 | asl |
| groupVII | 20101469 | nbeaa |
| groupVII | 20645749 | tiam1a |
| groupVII | 20655065 | tiam1a |
| groupVII | 22692904 | lrch2 |
| groupVII | 24337667 | zbtb16a |
| groupVII | 24480979 | ncam1a |
| groupVII | 25566233 | wwc1 |
| groupVII | 26822186 | doc2b |
| groupVII | 26867183 | rcc1l |
| groupVII | 27872941 | tmprss15 |
| groupVII | 27872950 | tmprss15 |
| groupVII | 27872952 | tmprss15 |
| groupVIII | 1029214 | nfia |
| groupVIII | 1067947 | nfia |
| groupVIII | 1255699 | sgta |
| groupVIII | 1261461 | SLC39A3 |
| groupVIII | 1444901 | slc1a7b |
| groupVIII | 1914639 | si:ch73-194h10.2 |
| groupVIII | 2065096 | lpp |
| groupVIII | 2119853 | magoh |
| groupVIII | 2498771 | tgfbr3 |
| groupVIII | 3513731 | irf4b |
| groupVIII | 3717572 | GMDS |
| groupVIII | 3729233 | GMDS |
| groupVIII | 4509668 | gpx7 |
| groupVIII | 4570277 | atpaf1 |
| groupVIII | 6561046 | meltf |
| groupVIII | 6684784 | slc25a24 |
| groupVIII | 7946080 | raver2 |
| groupVIII | 10306874 | ptprsa |
| groupVIII | 10310570 | ptprsa |
| groupVIII | 10550808 | nr5a2 |
| groupVIII | 11785209 | ankrd13c |
| groupVIII | 11795426 | cthl |
| groupVIII | 11925305 | negr1 |
| groupVIII | 12630691 | actl6a |
| groupVIII | 12866299 | si:dkey-30h22.11 |
| groupVIII | 12866318 | si:dkey-30h22.11 |
| groupVIII | 13723230 | cacna1ea |
| groupVIII | 14636941 | tle2b |
| groupVIII | 15002573 | si:dkey-97o5.1 |
| groupVIII | 15101479 | olfml2ba |
| groupVIII | 15101480 | olfml2ba |
| groupVIII | 15101511 | olfml2ba |
| groupVIII | 15136378 | GHRHR |
| groupVIII | 16815001 | si:ch211-212d10.2 |
| groupVIII | 16933459 | tmem161a |
| groupVIII | 17159656 | tjp3 |
| groupVIII | 17449991 | dph5 |
| groupVIII | 17820046 | gabrb3 |
| groupVIII | 17962542 | cyfip1 |
| groupVIII | 18044970 | tbc1d23 |
| groupVIII | 18497497 | sgcb |
| groupVIII | 18542341 | scfd2 |
| groupVIII | 18970188 | fsd1 |
| groupVIII | 19045555 | ddr2a |
| groupVIII | 19062127 | serbp1a |
| groupVIII | 19218952 | ralgps2 |
| groupVIII | 19247716 | ralgps2 |
| groupX | 1258350 | bmp8a |
| groupX | 1431303 | rnf19b |
| groupX | 1632737 | DPY19L4 |
| groupX | 1635764 | DPY19L4 |
| groupX | 1872357 | hivep1 |
| groupX | 1872395 | hivep1 |
| groupX | 2096192 | slc9a3.2 |
| groupX | 2651577 | tfap2e |
| groupX | 3893681 | STT3B |
| groupX | 4885550 | mylipa |
| groupX | 5119835 | bloc1s4 |
| groupX | 7976349 | mterf3 |
| groupX | 8921684 | calcr |
| groupX | 9043725 | cdk6 |
| groupX | 9136826 | TMEM196 |
| groupX | 9222085 | sp4 |
| groupX | 9263082 | col1a2 |
| groupX | 9705450 | rftn1a |
| groupX | 11348351 | fhod3a |
| groupX | 11365416 | fhod3a |
| groupX | 13560411 | TRAPPC9 |
| groupX | 14515807 | ccdc106b |
| groupX | 14675401 | rnf139 |
| groupX | 15588482 | bop1 |
| groupXI | 44440 | pdxdc1 |
| groupXI | 44441 | pdxdc1 |
| groupXI | 299682 | mettl22 |
| groupXI | 437592 | prrg2 |
| groupXI | 764525 | nags |
| groupXI | 1382735 | carm1 |
| groupXI | 1387661 | carm1 |
| groupXI | 2175473 | PSME3 |
| groupXI | 2313798 | caskin1 |
| groupXI | 3571147 | ntn1b |
| groupXI | 3974501 | GRIN2B |
| groupXI | 5013362 | TST |
| groupXI | 6507868 | st8sia6 |
| groupXI | 8663375 | ttyh2l |
| groupXI | 8717546 | aatka |
| groupXI | 8717548 | aatka |
| groupXI | 8726277 | aatka |
| groupXI | 10003166 | ADCY9 |
| groupXI | 10098607 | prkar1b |
| groupXI | 10817841 | arhgap17b |
| groupXI | 10935631 | noxo1a |
| groupXI | 10966681 | srebf1 |
| groupXI | 11449609 | 09-Sep |
| groupXI | 11519770 | tnrc6c1 |
| groupXI | 12093819 | gtpbp1 |
| groupXI | 12093831 | gtpbp1 |
| groupXI | 12656280 | mapk8ip3 |
| groupXI | 14254415 | si:dkey-43p13.5 |
| groupXI | 15562330 | fam83fa |
| groupXI | 15612231 | si:ch73-233k15.2 |
| groupXI | 15612252 | si:ch73-233k15.2 |
| groupXI | 15612253 | si:ch73-233k15.2 |
| groupXI | 15612265 | si:ch73-233k15.2 |
| groupXI | 15821746 | med15 |
| groupXI | 16487641 | tmem104 |
| groupXII | 1594422 | scube3 |
| groupXII | 1594424 | scube3 |
| groupXII | 2621029 | tm9sf4 |
| groupXII | 2621171 | tm9sf4 |
| groupXII | 3562883 | usp19 |
| groupXII | 3562888 | usp19 |
| groupXII | 4060908 | ampd1 |
| groupXII | 4060911 | ampd1 |
| groupXII | 4306905 | VSTM2L |
| groupXII | 5661366 | mib2 |
| groupXII | 5661843 | mib2 |
| groupXII | 5839118 | epha2b |
| groupXII | 5846364 | epha2b |
| groupXII | 5846559 | epha2b |
| groupXII | 7018377 | celsr3 |
| groupXII | 7100749 | mgll |
| groupXII | 7471705 | CACNA1D |
| groupXII | 7511555 | nsun5 |
| groupXII | 8433707 | KAZN |
| groupXII | 8835052 | phactr3a |
| groupXII | 9996128 | GSTM4 |
| groupXII | 11523684 | rarga |
| groupXII | 13531664 | slc16a1b |
| groupXII | 14077868 | plxna2 |
| groupXII | 14154789 | plxna2 |
| groupXII | 14268544 | magi3a |
| groupXII | 15489921 | pip4k2ca |
| groupXII | 15823000 | opn7d |
| groupXII | 15935495 | magixb |
| groupXII | 18272380 | rtel1 |
| groupXIII | 302905 | nelfb |
| groupXIII | 1762349 | hic2 |
| groupXIII | 2959586 | VPS37B |
| groupXIII | 3787612 | zmat4a |
| groupXIII | 4379022 | msh3 |
| groupXIII | 4549297 | lhfpl2b |
| groupXIII | 7190703 | frmd3 |
| groupXIII | 8461578 | ebf2 |
| groupXIII | 10266430 | cox7c |
| groupXIII | 11442150 | adgrv1 |
| groupXIII | 12071325 | bmp1a |
| groupXIII | 12116272 | gfpt1 |
| groupXIII | 12139588 | aak1a |
| groupXIII | 12392757 | tcf7l1a |
| groupXIII | 12863489 | rph3ab |
| groupXIII | 13129213 | trpm6 |
| groupXIII | 13182723 | grk5l |
| groupXIII | 13185867 | grk5l |
| groupXIII | 13185885 | grk5l |
| groupXIII | 13185886 | grk5l |
| groupXIII | 14354401 | ncor2 |
| groupXIII | 14850519 | kcnv2b |
| groupXIII | 16347129 | specc1la |
| groupXIII | 16372968 | si:dkey-91m11.5 |
| groupXIII | 17236935 | jak2b |
| groupXIII | 17405512 | poli |
| groupXIII | 17568669 | notch1b |
| groupXIII | 17736887 | col27a1b |
| groupXIII | 19303002 | fbp2 |
| groupXIII | 19642070 | coq5 |
| groupXIV | 549351 | ZSWIM6 |
| groupXIV | 808642 | col4a3bpb |
| groupXIV | 916126 | wdr41 |
| groupXIV | 1031933 | ARSB |
| groupXIV | 1405959 | fcho2 |
| groupXIV | 2538572 | aif1l |
| groupXIV | 2732354 | kyat1 |
| groupXIV | 2780479 | NTNG2 |
| groupXIV | 4411253 | TMEM38B |
| groupXIV | 4960232 | ASTN2 |
| groupXIV | 7011755 | KCNN2 |
| groupXIV | 7375043 | morn5 |
| groupXIV | 7925226 | cdc37l1 |
| groupXIV | 8130524 | bcr |
| groupXIV | 8307713 | si:dkey-112m2.1 |
| groupXIV | 8341222 | si:dkey-112m2.1 |
| groupXIV | 8612841 | sfswap |
| groupXIV | 9394041 | plppr1 |
| groupXIV | 9394042 | plppr1 |
| groupXIV | 9403163 | plppr1 |
| groupXIV | 10153615 | rab27b |
| groupXIV | 13361141 | spina |
| groupXIV | 13494921 | dcc |
| groupXIV | 13541022 | dcc |
| groupXIV | 13802010 | CRB2 |
| groupXIV | 14608983 | ntmt1 |
| groupXIV | 15077079 | auh |
| groupXIX | 279445 | ntrk3a |
| groupXIX | 309244 | ntrk3a |
| groupXIX | 315383 | ntrk3a |
| groupXIX | 815471 | ndrg4 |
| groupXIX | 1002818 | cry2 |
| groupXIX | 1002823 | cry2 |
| groupXIX | 1002824 | cry2 |
| groupXIX | 1002836 | cry2 |
| groupXIX | 1149661 | galnt18a |
| groupXIX | 1240432 | PPFIBP2 |
| groupXIX | 1284472 | far1 |
| groupXIX | 1485001 | ric8a |
| groupXIX | 1580409 | zgc:91860 |
| groupXIX | 1607502 | ces3 |
| groupXIX | 2123911 | lonp2 |
| groupXIX | 2216564 | st3gal2 |
| groupXIX | 2259685 | hist1h4l |
| groupXIX | 3055852 | klhdc4 |
| groupXIX | 3343656 | kiaa1549la |
| groupXIX | 5448302 | nell2a |
| groupXIX | 6275171 | usp3 |
| groupXIX | 6306183 | commd4 |
| groupXIX | 6646006 | smad3b |
| groupXIX | 7581386 | tjp1b |
| groupXIX | 7799647 | INSC |
| groupXIX | 8132582 | hsd17b12a |
| groupXIX | 8586228 | ptpn9a |
| groupXIX | 10488868 | si:dkey-106n21.1 |
| groupXIX | 10919586 | lrrk2 |
| groupXIX | 11256691 | tspan18b |
| groupXIX | 11744277 | PRKAR2B |
| groupXIX | 12704045 | si:ch211-13k12.2 |
| groupXIX | 12704047 | si:ch211-13k12.2 |
| groupXIX | 13112677 | dusp8a |
| groupXIX | 13808858 | rassf7a |
| groupXIX | 15891442 | shank3a |
| groupXIX | 17610748 | uri1 |
| groupXIX | 17610751 | uri1 |
| groupXIX | 17610753 | uri1 |
| groupXIX | 17610757 | uri1 |
| groupXIX | 18253089 | mical3a |
| groupXIX | 18568168 | tmtc2b |
| groupXIX | 18720711 | nup205 |
| groupXIX | 18907475 | kcng4a |
| groupXV | 141476 | arid4a |
| groupXV | 187245 | BAZ1A |
| groupXV | 352022 | atrn |
| groupXV | 361817 | atrn |
| groupXV | 391375 | eif2ak3 |
| groupXV | 923132 | klc1a |
| groupXV | 986791 | BRF1 |
| groupXV | 994919 | BRF1 |
| groupXV | 1329085 | mrps26 |
| groupXV | 1397019 | slc39a8 |
| groupXV | 2369927 | RYR3 |
| groupXV | 2915763 | rbbp4 |
| groupXV | 4126558 | pabpc4 |
| groupXV | 5506022 | hspa4l |
| groupXV | 5882707 | eml1 |
| groupXV | 6034997 | rpap1 |
| groupXV | 7048473 | si:ch211-15d5.11 |
| groupXV | 7048495 | si:ch211-15d5.11 |
| groupXV | 8912484 | fermt2 |
| groupXV | 8952097 | STXBP6 |
| groupXV | 10021510 | fndc3ba |
| groupXV | 10561857 | elmsan1b |
| groupXV | 10874243 | slc25a21 |
| groupXV | 11634105 | sos2 |
| groupXV | 11928511 | pacs2 |
| groupXV | 12056687 | slc8a3 |
| groupXV | 12570065 | lrfn5a |
| groupXV | 15509173 | gabrb1 |
| groupXV | 16002575 | iars2 |
| groupXVI | 147492 | slc37a1 |
| groupXVI | 147494 | slc37a1 |
| groupXVI | 195347 | pde9a |
| groupXVI | 1629241 | clasp1a |
| groupXVI | 2474886 | mycbp2 |
| groupXVI | 3030302 | nalcn |
| groupXVI | 4490055 | asb1 |
| groupXVI | 4549159 | hdac4 |
| groupXVI | 4565118 | hdac4 |
| groupXVI | 5755922 | gulp1a |
| groupXVI | 6394706 | glsb |
| groupXVI | 6583007 | IMPG2 |
| groupXVI | 6621698 | abi3bpa |
| groupXVI | 6622524 | abi3bpa |
| groupXVI | 6645406 | tfg |
| groupXVI | 7726845 | appb |
| groupXVI | 9603361 | prkra |
| groupXVI | 10223788 | SLC25A12 |
| groupXVI | 10554458 | stk39 |
| groupXVI | 11685983 | nme8 |
| groupXVI | 11686070 | nme8 |
| groupXVI | 11696630 | SMARCAL1 |
| groupXVI | 11739083 | pfkla |
| groupXVI | 11741264 | pfkla |
| groupXVI | 11799716 | trpm2 |
| groupXVI | 12337419 | vwc2l |
| groupXVI | 13042576 | SH3RF3 |
| groupXVI | 15761756 | ABI2 |
| groupXVI | 16555594 | abcb6b |
| groupXVI | 16624511 | kalrna |
| groupXVI | 16910614 | parp4 |
| groupXVI | 17292751 | itgb5 |
| groupXVI | 17347979 | dgkg |
| groupXVI | 17462557 | pikfyve |
| groupXVI | 17685457 | rab5b |
| groupXVI | 18091809 | mao |
| groupXVII | 486408 | ctsa |
| groupXVII | 558224 | mkrn2 |
| groupXVII | 833691 | dnase1l4.1 |
| groupXVII | 1664612 | atp2b2 |
| groupXVII | 1978579 | arhgef10la |
| groupXVII | 2999932 | pacsin1b |
| groupXVII | 3506319 | slc5a8l |
| groupXVII | 4011290 | ASIC1 |
| groupXVII | 4041973 | ASIC1 |
| groupXVII | 4183577 | eya2 |
| groupXVII | 4366611 | cdh22 |
| groupXVII | 4759550 | st7l |
| groupXVII | 5555572 | MAP3K12 |
| groupXVII | 6549208 | cpne5b |
| groupXVII | 7158503 | krt4 |
| groupXVII | 10275514 | dido1 |
| groupXVII | 10413238 | zgc:92107 |
| groupXVII | 11027907 | txnrd3 |
| groupXVII | 11255412 | etnk2 |
| groupXVII | 11308336 | klhdc8a |
| groupXVII | 11955141 | wnt7aa |
| groupXVII | 13238989 | EPB41L1 |
| groupXVII | 14069629 | mitfa |
| groupXVII | 14215418 | itpr1b |
| groupXVII | 14249407 | arl8bb |
| groupXVIII | 293689 | lbr |
| groupXVIII | 496239 | cnih3 |
| groupXVIII | 631355 | ptchd4 |
| groupXVIII | 639121 | ptchd4 |
| groupXVIII | 2332942 | nrxn3b |
| groupXVIII | 2394844 | nrxn3b |
| groupXVIII | 2544694 | cyp46a1.4 |
| groupXVIII | 3033960 | mcm9 |
| groupXVIII | 3446131 | mdn1 |
| groupXVIII | 4499462 | si:dkey-119m7.4 |
| groupXVIII | 4499526 | si:dkey-119m7.4 |
| groupXVIII | 7671230 | slc24a4b |
| groupXVIII | 8566642 | afg1lb |
| groupXVIII | 8579462 | afg1lb |
| groupXVIII | 9543664 | rhag |
| groupXVIII | 9763683 | slc35f6 |
| groupXVIII | 10240054 | daam1b |
| groupXVIII | 10241528 | daam1b |
| groupXVIII | 10843673 | pinx1 |
| groupXVIII | 11889168 | ust |
| groupXVIII | 12136724 | stxbp5a |
| groupXVIII | 12179367 | rab32a |
| groupXVIII | 12721986 | myt1la |
| groupXVIII | 13771632 | wdcp |
| groupXX | 85188 | dennd4b |
| groupXX | 172527 | pik3r4 |
| groupXX | 633589 | tmem145 |
| groupXX | 1984914 | snrka |
| groupXX | 2463673 | spire1b |
| groupXX | 2557791 | ubr5 |
| groupXX | 2614088 | oxr1b |
| groupXX | 2705937 | si:dkey-122a22.2 |
| groupXX | 3669526 | gdf6a |
| groupXX | 3669532 | gdf6a |
| groupXX | 3669614 | gdf6a |
| groupXX | 4980646 | derl1 |
| groupXX | 5025141 | atad2 |
| groupXX | 5025181 | atad2 |
| groupXX | 5036765 | psma2 |
| groupXX | 6155595 | RBMS3 |
| groupXX | 6965505 | nrsn1 |
| groupXX | 7033062 | nrsn1 |
| groupXX | 7048536 | nrsn1 |
| groupXX | 8808003 | si:ch73-380l3.2 |
| groupXX | 10712460 | rad54b |
| groupXX | 10824480 | fam171a1 |
| groupXX | 12961899 | kcnn3 |
| groupXX | 13183746 | flad1 |
| groupXX | 14703868 | VWDE |
| groupXX | 15626825 | smg5 |
| groupXX | 15626826 | smg5 |
| groupXX | 15626830 | smg5 |
| groupXX | 15626833 | smg5 |
| groupXX | 16270708 | onecutl |
| groupXX | 17274290 | si:ch211-1i11.3 |
| groupXX | 18131353 | tgfb2 |
| groupXXI | 1091376 | PARD3 |
| groupXXI | 3762843 | fen1 |
| groupXXI | 3766074 | fen1 |
| groupXXI | 4579413 | dap |
| groupXXI | 5686739 | mllt10 |
| groupXXI | 5966696 | pcyt1ba |
| groupXXI | 6708043 | KCNB2 |
| groupXXI | 7549622 | pan3 |
| groupXXI | 7879936 | vipr1b |
| groupXXI | 8687826 | acad11 |
| groupXXI | 8829563 | dlgap1b |
| groupXXI | 8928532 | insig1 |
| groupXXI | 9261362 | esyt2b |
| groupXXI | 9261363 | esyt2b |
| groupXXI | 9261365 | esyt2b |
| groupXXI | 9261366 | esyt2b |
| groupXXI | 10219841 | DNAJC13 |
| groupXXI | 11076874 | gpr158a |
