## Supplemental Table 5 for "Using seasonal genomic changes to understand historical adaptation: parallel selection on stickleback in highly-variable estuaries"

| Table S5. F_ST_-qGLM outliers that map to protein-coding genes. | | |
| --- | --- | --- |
| Chromosome | Position | Overlapping gene |
| groupI | 1356415 | myo1cb |
| groupI | 1356417 | myo1cb |
| groupI | 1356419 | myo1cb |
| groupI | 2225425 | CEP164 |
| groupI | 2250676 | sik3 |
| groupI | 11997404 | pitpnm3 |
| groupI | 15136838 | nectin1b |
| groupI | 18856683 | bcas3 |
| groupI | 20075909 | bmpr2a |
| groupI | 22824368 | cacnb4b |
| groupI | 23628696 | GABRG3 |
| groupI | 27700431 | cybb |
| groupII | 267489 | aldh1a3 |
| groupII | 2023253 | si:ch211-186j3.6 |
| groupII | 5046745 | kif13ba |
| groupII | 6086839 | pir |
| groupII | 7214728 | diaph3 |
| groupII | 9373814 | ext2 |
| groupII | 9489069 | plekha7b |
| groupII | 10284123 | galnt18b |
| groupII | 10284126 | galnt18b |
| groupII | 11413867 | cgnl1 |
| groupII | 13419593 | supv3l1 |
| groupII | 19363379 | MYO9A |
| groupII | 19373274 | MYO9A |
| groupII | 19529077 | neo1b |
| groupII | 21682346 | si:ch211-112g6.4 |
| groupIII | 4120217 | ctnnd2b |
| groupIII | 5202005 | aglb |
| groupIII | 5593347 | sdr16c5a |
| groupIII | 6977810 | wdr47a |
| groupIII | 12400167 | insra |
| groupIII | 16578504 | AGAP3 |
| groupIV | 2063943 | si:dkey-237h12.3 |
| groupIV | 2945152 | LIMCH1 |
| groupIV | 3525588 | ANKRD50 |
| groupIV | 7963436 | BEND4 |
| groupIV | 18340744 | grm8a |
| groupIV | 29377073 | chchd3a |
| groupIV | 30675359 | FUT9 |
| groupIV | 30675360 | FUT9 |
| groupIV | 32062097 | btbd11a |
| groupIV | 32330935 | ERI1 |
| groupIX | 170530 | adgrl3.1 |
| groupIX | 2282295 | SLC24A3 |
| groupIX | 5374892 | trim2a |
| groupIX | 6047838 | anxa5b |
| groupIX | 6047839 | anxa5b |
| groupIX | 6789945 | pou4f2 |
| groupIX | 7554047 | sh3gl2a |
| groupIX | 18849813 | strn |
| groupIX | 20141599 | zgc:109889 |
| groupV | 3530315 | TANC2 |
| groupV | 5771266 | rangap1b |
| groupV | 6301445 | fdxr |
| groupVI | 65315 | pdzd8 |
| groupVI | 4144174 | smap1 |
| groupVI | 4146999 | smap1 |
| groupVI | 4147000 | smap1 |
| groupVI | 5854986 | rrbp1a |
| groupVI | 10098698 | kcnma1a |
| groupVI | 10512756 | polh |
| groupVI | 10695465 | pcdh15a |
| groupVI | 14491371 | SPTBN1 |
| groupVI | 16698814 | entpd6 |
| groupVI | 16724969 | ninl |
| groupVII | 734163 | hdac8 |
| groupVII | 2070514 | arhgef11 |
| groupVII | 2281161 | pacs1a |
| groupVII | 2297755 | pacs1a |
| groupVII | 10336150 | senp3b |
| groupVII | 14493165 | p2rx1 |
| groupVII | 16078035 | MSI2 |
| groupVII | 18430464 | grk1b |
| groupVII | 21446485 | slc47a1 |
| groupVII | 21450557 | slc47a1 |
| groupVII | 21464005 | slc47a1 |
| groupVII | 26806302 | doc2b |
| groupVII | 27094361 | pou2f3 |
| groupVII | 27182881 | slc22a5 |
| groupVIII | 1025715 | nfia |
| groupVIII | 1235981 | nwd1 |
| groupVIII | 1252454 | thop1 |
| groupVIII | 1261437 | SLC39A3 |
| groupVIII | 1261461 | SLC39A3 |
| groupVIII | 1446795 | slc1a7b |
| groupVIII | 1914639 | si:ch73-194h10.2 |
| groupVIII | 6344609 | mccc1 |
| groupVIII | 7117808 | znhit6 |
| groupVIII | 10330503 | tbxa2r |
| groupVIII | 13898442 | faf1 |
| groupVIII | 14408125 | lpl |
| groupVIII | 17323822 | grin3bb |
| groupVIII | 17357959 | med16 |
| groupVIII | 17357961 | med16 |
| groupVIII | 17820046 | gabrb3 |
| groupVIII | 18066094 | sft2d2a |
| groupVIII | 18605846 | lnx1 |
| groupX | 1265572 | MACF1 |
| groupX | 6434251 | AKAP9 |
| groupX | 7428667 | KCNQ4 |
| groupX | 7796551 | spire1a |
| groupX | 9712971 | rftn1a |
| groupX | 13158201 | ST3GAL1 |
| groupX | 13612904 | TRAPPC9 |
| groupX | 14529968 | cnot3b |
| groupX | 15057778 | vps52 |
| groupXI | 4768115 | smcr8a |
| groupXI | 9089704 | cyth1a |
| groupXI | 9093791 | cyth1a |
| groupXI | 11460524 | 09-Sep |
| groupXI | 13038226 | akt1s1 |
| groupXI | 14802584 | tnrc6a |
| groupXI | 14903507 | prkcbb |
| groupXII | 334413 | dnah12 |
| groupXII | 1406570 | g6pd |
| groupXII | 1439605 | asic1a |
| groupXII | 7474444 | CACNA1D |
| groupXII | 13564965 | lima1a |
| groupXII | 15674173 | osbpl2a |
| groupXIII | 277188 | ehmt1a |
| groupXIII | 6275830 | elovl7a |
| groupXIII | 8091956 | pcsk5b |
| groupXIII | 8583186 | tcn2 |
| groupXIII | 13395404 | DTX1 |
| groupXIII | 13888895 | zgc:154046 |
| groupXIII | 18809089 | wdfy3 |
| groupXIII | 19100744 | adamts3 |
| groupXIII | 19368639 | atp8b5a |
| groupXIV | 916126 | wdr41 |
| groupXIV | 7724003 | dbh |
| groupXIV | 7925224 | cdc37l1 |
| groupXIV | 7925227 | cdc37l1 |
| groupXIV | 12051392 | rxraa |
| groupXIV | 14709730 | YTHDC2 |
| groupXIV | 14854047 | slc38a9 |
| groupXIX | 702930 | exoc3l1 |
| groupXIX | 758493 | smpd3 |
| groupXIX | 1259055 | kdelc2 |
| groupXIX | 1481801 | ric8a |
| groupXIX | 2822284 | nrn1la |
| groupXIX | 6006011 | zgc:136858 |
| groupXIX | 6773313 | tle3b |
| groupXIX | 7058825 | pgghg |
| groupXIX | 7432798 | MYO5A |
| groupXIX | 7781050 | INSC |
| groupXIX | 7803011 | INSC |
| groupXIX | 10201370 | cftr |
| groupXIX | 10225694 | asz1 |
| groupXIX | 10225695 | asz1 |
| groupXIX | 11438458 | RELN |
| groupXIX | 13808858 | rassf7a |
| groupXIX | 14035769 | pamr1 |
| groupXIX | 14035773 | pamr1 |
| groupXIX | 14073941 | slc1a2b |
| groupXIX | 16675756 | lmf2a |
| groupXIX | 17562354 | srgap1a |
| groupXV | 56921 | mapkbp1 |
| groupXV | 985430 | BRF1 |
| groupXV | 5388528 | OTUB2 |
| groupXV | 10244764 | grin3ba |
| groupXV | 13281395 | pdss2 |
| groupXVI | 3059660 | nalcn |
| groupXVI | 3069359 | ITGBL1 |
| groupXVI | 3209162 | p2ry8 |
| groupXVI | 6455500 | slc40a1 |
| groupXVI | 10222778 | SLC25A12 |
| groupXVI | 12337419 | vwc2l |
| groupXVII | 5344466 | ARHGAP9 |
| groupXVII | 11237013 | sox13 |
| groupXVIII | 747441 | tnfrsf21 |
| groupXVIII | 10843666 | pinx1 |
| groupXVIII | 11827808 | METTL24 |
| groupXVIII | 12137126 | stxbp5a |
| groupXVIII | 13768203 | fkbp1b |
| groupXVIII | 14165771 | PLCB1 |
| groupXX | 518577 | si:dkey-23o4.6 |
| groupXX | 6575071 | adamts16 |
| groupXX | 6934056 | nrsn1 |
| groupXX | 7484682 | cdkal1 |
| groupXX | 8793862 | si:ch73-380l3.2 |
| groupXX | 8814434 | si:ch73-380l3.2 |
| groupXX | 10710861 | rad54b |
| groupXX | 10865868 | abcb4 |
| groupXX | 16253625 | ctss2.1 |
| groupXX | 18939681 | cdk14 |
| groupXX | 18957089 | cdk14 |
| groupXX | 18957777 | cdk14 |
| groupXXI | 1068422 | PARD3 |
| groupXXI | 9447756 | SLC39A12 |
