## Supplemental Table 6 for "Using seasonal genomic changes to understand historical adaptation: parallel selection on stickleback in highly-variable estuaries"

| Table S6. Enrichment of molecular functions among F_ST_ outlier genes (P-value < 0.05). | | | | | |
| --- | --- | --- | --- | --- | --- |
| GO.ID | Term | Annotated | Significant | Expected | classic |
| GO:0005509 | calcium ion binding | 407 | 39 | 18.87 | 1.20E-05 |
| GO:0005201 | extracellular matrix structural constitu... | 23 | 6 | 1.07 | 0.00049 |
| GO:0005515 | protein binding | 3356 | 187 | 155.62 | 0.00101 |
| GO:0043167 | ion binding | 3318 | 181 | 153.85 | 0.00374 |
| GO:0043169 | cation binding | 1937 | 112 | 89.82 | 0.00528 |
| GO:0046872 | metal ion binding | 1924 | 111 | 89.22 | 0.00591 |
| GO:0015085 | calcium ion transmembrane transporter ac... | 74 | 9 | 3.43 | 0.00695 |
| GO:0005488 | binding | 7189 | 355 | 333.35 | 0.0101 |
| GO:0004000 | adenosine deaminase activity | 11 | 3 | 0.51 | 0.01237 |
| GO:0003779 | actin binding | 190 | 16 | 8.81 | 0.01523 |
| GO:0015368 | calcium:cation antiporter activity | 12 | 3 | 0.56 | 0.01593 |
| GO:0008092 | cytoskeletal protein binding | 307 | 22 | 14.24 | 0.02813 |
| GO:0005096 | GTPase activator activity | 109 | 10 | 5.05 | 0.02958 |
| GO:0003774 | motor activity | 94 | 9 | 4.36 | 0.02988 |
| GO:0016814 | hydrolase activity, acting on carbon-nit... | 26 | 4 | 1.21 | 0.03035 |
| GO:0030695 | GTPase regulator activity | 110 | 10 | 5.1 | 0.03125 |
| GO:0016462 | pyrophosphatase activity | 513 | 33 | 23.79 | 0.03463 |
| GO:0016887 | ATPase activity | 112 | 10 | 5.19 | 0.03479 |
| GO:0016817 | hydrolase activity, acting on acid anhyd... | 515 | 33 | 23.88 | 0.03629 |
| GO:0016818 | hydrolase activity, acting on acid anhyd... | 515 | 33 | 23.88 | 0.03629 |
| GO:0008081 | phosphoric diester hydrolase activity | 68 | 7 | 3.15 | 0.03738 |
| GO:0032559 | adenyl ribonucleotide binding | 1095 | 63 | 50.77 | 0.03985 |
| GO:0030554 | adenyl nucleotide binding | 1097 | 63 | 50.87 | 0.04112 |
| GO:0005524 | ATP binding | 1079 | 62 | 50.03 | 0.0422 |
| GO:0060589 | nucleoside-triphosphatase regulator acti... | 116 | 10 | 5.38 | 0.04268 |
| GO:0016409 | palmitoyltransferase activity | 29 | 4 | 1.34 | 0.04328 |
| GO:0005544 | calcium-dependent phospholipid binding | 18 | 3 | 0.83 | 0.04813 |
| GO:0016810 | hydrolase activity, acting on carbon-nit... | 72 | 7 | 3.34 | 0.04863 |
