## Supplemental Table 7 for "Using seasonal genomic changes to understand historical adaptation: parallel selection on stickleback in highly-variable estuaries"

| Table S7. Enrichment of molecular functions of genes harbouring qGLM outliers (P-value < 0.05). | | | | | |
| --- | --- | --- | --- | --- | --- |
| GO.ID | Term | Annotated | Significant | Expected | classic |
| GO:0046873 | metal ion transmembrane transporter acti... | 247 | 29 | 11.13 | 2.00E-06 |
| GO:0015075 | ion transmembrane transporter activity | 506 | 46 | 22.79 | 3.70E-06 |
| GO:0015085 | calcium ion transmembrane transporter ac... | 74 | 14 | 3.33 | 4.40E-06 |
| GO:0005216 | ion channel activity | 281 | 29 | 12.66 | 2.50E-05 |
| GO:0022838 | substrate-specific channel activity | 283 | 29 | 12.75 | 2.90E-05 |
| GO:0022890 | inorganic cation transmembrane transport... | 331 | 32 | 14.91 | 3.60E-05 |
| GO:0008324 | cation transmembrane transporter activit... | 350 | 33 | 15.77 | 4.50E-05 |
| GO:0015318 | inorganic molecular entity transmembrane... | 461 | 40 | 20.77 | 4.80E-05 |
| GO:0005261 | cation channel activity | 198 | 22 | 8.92 | 8.10E-05 |
| GO:0015267 | channel activity | 305 | 29 | 13.74 | 0.00011 |
| GO:0022803 | passive transmembrane transporter activi... | 305 | 29 | 13.74 | 0.00011 |
| GO:0022839 | ion gated channel activity | 236 | 24 | 10.63 | 0.00016 |
| GO:0005262 | calcium channel activity | 56 | 10 | 2.52 | 0.00017 |
| GO:0005515 | protein binding | 3356 | 187 | 151.18 | 0.00018 |
| GO:0022836 | gated channel activity | 241 | 24 | 10.86 | 0.00022 |
| GO:0005516 | calmodulin binding | 51 | 9 | 2.3 | 0.0004 |
| GO:0019899 | enzyme binding | 237 | 22 | 10.68 | 0.00102 |
| GO:0022857 | transmembrane transporter activity | 748 | 52 | 33.7 | 0.00103 |
| GO:0005215 | transporter activity | 771 | 53 | 34.73 | 0.00119 |
| GO:0051020 | GTPase binding | 189 | 17 | 8.51 | 0.00508 |
| GO:0017048 | Rho GTPase binding | 26 | 5 | 1.17 | 0.00544 |
| GO:0005488 | binding | 7189 | 347 | 323.85 | 0.00573 |
| GO:0015079 | potassium ion transmembrane transporter ... | 102 | 11 | 4.59 | 0.00606 |
| GO:0005085 | guanyl-nucleotide exchange factor activi... | 133 | 13 | 5.99 | 0.00687 |
| GO:0004709 | MAP kinase kinase kinase activity | 10 | 3 | 0.45 | 0.0086 |
| GO:0005347 | ATP transmembrane transporter activity | 29 | 5 | 1.31 | 0.00879 |
| GO:0022843 | voltage-gated cation channel activity | 93 | 10 | 4.19 | 0.00886 |
| GO:0000295 | adenine nucleotide transmembrane transpo... | 30 | 5 | 1.35 | 0.01017 |
| GO:0005346 | purine ribonucleotide transmembrane tran... | 30 | 5 | 1.35 | 0.01017 |
| GO:0015215 | nucleotide transmembrane transporter act... | 30 | 5 | 1.35 | 0.01017 |
| GO:0015216 | purine nucleotide transmembrane transpor... | 30 | 5 | 1.35 | 0.01017 |
| GO:0015605 | organophosphate ester transmembrane tran... | 30 | 5 | 1.35 | 0.01017 |
| GO:0015491 | cation:cation antiporter activity | 20 | 4 | 0.9 | 0.01108 |
| GO:0005227 | calcium activated cation channel activit... | 11 | 3 | 0.5 | 0.01143 |
| GO:0015278 | calcium-release channel activity | 11 | 3 | 0.5 | 0.01143 |
| GO:0099604 | ligand-gated calcium channel activity | 11 | 3 | 0.5 | 0.01143 |
| GO:0015298 | solute:cation antiporter activity | 21 | 4 | 0.95 | 0.01321 |
| GO:0015276 | ligand-gated ion channel activity | 114 | 11 | 5.14 | 0.01356 |
| GO:0022834 | ligand-gated channel activity | 114 | 11 | 5.14 | 0.01356 |
| GO:0015368 | calcium:cation antiporter activity | 12 | 3 | 0.54 | 0.01474 |
| GO:0005245 | voltage-gated calcium channel activity | 22 | 4 | 0.99 | 0.01558 |
| GO:0043167 | ion binding | 3318 | 171 | 149.47 | 0.01594 |
| GO:0005509 | calcium ion binding | 407 | 28 | 18.33 | 0.0167 |
| GO:0015077 | monovalent inorganic cation transmembran... | 200 | 16 | 9.01 | 0.01851 |
| GO:0005088 | Ras guanyl-nucleotide exchange factor ac... | 13 | 3 | 0.59 | 0.01853 |
| GO:0004672 | protein kinase activity | 484 | 32 | 21.8 | 0.01856 |
| GO:0004674 | protein serine/threonine kinase activity | 251 | 19 | 11.31 | 0.0186 |
| GO:0016773 | phosphotransferase activity, alcohol gro... | 564 | 36 | 25.41 | 0.02122 |
| GO:0016301 | kinase activity | 604 | 38 | 27.21 | 0.02233 |
| GO:0005267 | potassium channel activity | 92 | 9 | 4.14 | 0.02244 |
| GO:0015238 | drug transmembrane transporter activity | 37 | 5 | 1.67 | 0.02408 |
| GO:0098772 | molecular function regulator | 525 | 33 | 23.65 | 0.03227 |
| GO:0005244 | voltage-gated ion channel activity | 130 | 11 | 5.86 | 0.03248 |
| GO:1901505 | carbohydrate derivative transmembrane tr... | 40 | 5 | 1.8 | 0.03259 |
| GO:0017016 | Ras GTPase binding | 68 | 7 | 3.06 | 0.0327 |
| GO:0031267 | small GTPase binding | 68 | 7 | 3.06 | 0.0327 |
| GO:0004890 | GABA-A receptor activity | 16 | 3 | 0.72 | 0.03283 |
| GO:0005217 | intracellular ligand-gated ion channel a... | 16 | 3 | 0.72 | 0.03283 |
| GO:0032559 | adenyl ribonucleotide binding | 1095 | 62 | 49.33 | 0.03285 |
| GO:0030554 | adenyl nucleotide binding | 1097 | 62 | 49.42 | 0.03393 |
| GO:0015932 | nucleobase-containing compound transmemb... | 41 | 5 | 1.85 | 0.0358 |
| GO:0043168 | anion binding | 1632 | 88 | 73.52 | 0.036 |
| GO:0022832 | voltage-gated channel activity | 134 | 11 | 6.04 | 0.03923 |
| GO:0008144 | drug binding | 1147 | 64 | 51.67 | 0.03944 |
| GO:0099516 | ion antiporter activity | 29 | 4 | 1.31 | 0.03956 |
| GO:0008514 | organic anion transmembrane transporter ... | 57 | 6 | 2.57 | 0.04224 |
| GO:0008195 | phosphatidate phosphatase activity | 18 | 3 | 0.81 | 0.04479 |
| GO:0016917 | GABA receptor activity | 18 | 3 | 0.81 | 0.04479 |
| GO:0004725 | protein tyrosine phosphatase activity | 73 | 7 | 3.29 | 0.0455 |
| GO:0005524 | ATP binding | 1079 | 60 | 48.61 | 0.04806 |
| GO:0008509 | anion transmembrane transporter activity | 139 | 11 | 6.26 | 0.04898 |
