## Supplemental Table 8 for "Using seasonal genomic changes to understand historical adaptation: parallel selection on stickleback in highly-variable estuaries"

| Table S8. Molecular functions enriched among F_ST_-qGLM genes (P-value < 0.05) | | | | | |
| --- | --- | --- | --- | --- | --- |
| GO.ID | Term | Annotated | Significant | Expected | classic |
| GO:0015075 | ion transmembrane transporter activity | 506 | 21 | 7 | 5.60E-06 |
| GO:0005215 | transporter activity | 771 | 25 | 10.66 | 4.80E-05 |
| GO:0015318 | inorganic molecular entity transmembrane... | 461 | 18 | 6.38 | 6.20E-05 |
| GO:0022857 | transmembrane transporter activity | 748 | 24 | 10.34 | 8.10E-05 |
| GO:0015276 | ligand-gated ion channel activity | 114 | 8 | 1.58 | 0.00017 |
| GO:0022834 | ligand-gated channel activity | 114 | 8 | 1.58 | 0.00017 |
| GO:0022839 | ion gated channel activity | 236 | 11 | 3.26 | 0.00042 |
| GO:0022836 | gated channel activity | 241 | 11 | 3.33 | 0.0005 |
| GO:0005216 | ion channel activity | 281 | 12 | 3.89 | 0.00051 |
| GO:0022838 | substrate-specific channel activity | 283 | 12 | 3.91 | 0.00054 |
| GO:0005230 | extracellular ligand-gated ion channel a... | 78 | 6 | 1.08 | 0.00071 |
| GO:0015267 | channel activity | 305 | 12 | 4.22 | 0.00104 |
| GO:0022803 | passive transmembrane transporter activi... | 305 | 12 | 4.22 | 0.00104 |
| GO:0008324 | cation transmembrane transporter activit... | 350 | 13 | 4.84 | 0.00109 |
| GO:0003779 | actin binding | 190 | 9 | 2.63 | 0.00126 |
| GO:0004890 | GABA-A receptor activity | 16 | 3 | 0.22 | 0.00127 |
| GO:0016917 | GABA receptor activity | 18 | 3 | 0.25 | 0.00181 |
| GO:0003774 | motor activity | 94 | 6 | 1.3 | 0.00188 |
| GO:0008509 | anion transmembrane transporter activity | 139 | 7 | 1.92 | 0.00311 |
| GO:0005254 | chloride channel activity | 49 | 4 | 0.68 | 0.00458 |
| GO:0022890 | inorganic cation transmembrane transport... | 331 | 11 | 4.58 | 0.00612 |
| GO:0005253 | anion channel activity | 55 | 4 | 0.76 | 0.00692 |
| GO:0046873 | metal ion transmembrane transporter acti... | 247 | 9 | 3.42 | 0.00724 |
| GO:0015108 | chloride transmembrane transporter activ... | 57 | 4 | 0.79 | 0.00785 |
| GO:0035381 | ATP-gated ion channel activity | 10 | 2 | 0.14 | 0.00794 |
| GO:0008092 | cytoskeletal protein binding | 307 | 10 | 4.25 | 0.01009 |
| GO:0015368 | calcium:cation antiporter activity | 12 | 2 | 0.17 | 0.01144 |
| GO:0005543 | phospholipid binding | 102 | 5 | 1.41 | 0.01344 |
| GO:0015297 | antiporter activity | 38 | 3 | 0.53 | 0.01534 |
| GO:0001614 | purinergic nucleotide receptor activity | 14 | 2 | 0.19 | 0.0155 |
| GO:0016502 | nucleotide receptor activity | 14 | 2 | 0.19 | 0.0155 |
| GO:0005096 | GTPase activator activity | 109 | 5 | 1.51 | 0.01747 |
| GO:0030695 | GTPase regulator activity | 110 | 5 | 1.52 | 0.0181 |
| GO:0008289 | lipid binding | 202 | 7 | 2.79 | 0.02195 |
| GO:0060589 | nucleoside-triphosphatase regulator acti... | 116 | 5 | 1.6 | 0.02223 |
| GO:0005544 | calcium-dependent phospholipid binding | 18 | 2 | 0.25 | 0.02513 |
| GO:0015491 | cation:cation antiporter activity | 20 | 2 | 0.28 | 0.03066 |
| GO:0035586 | purinergic receptor activity | 20 | 2 | 0.28 | 0.03066 |
| GO:0015103 | inorganic anion transmembrane transporte... | 87 | 4 | 1.2 | 0.03227 |
| GO:0005516 | calmodulin binding | 51 | 3 | 0.71 | 0.03327 |
| GO:0015298 | solute:cation antiporter activity | 21 | 2 | 0.29 | 0.03358 |
| GO:0008047 | enzyme activator activity | 135 | 5 | 1.87 | 0.03915 |
| GO:0022804 | active transmembrane transporter activit... | 182 | 6 | 2.52 | 0.04044 |
| GO:0004970 | ionotropic glutamate receptor activity | 24 | 2 | 0.33 | 0.04297 |
| GO:0008066 | glutamate receptor activity | 24 | 2 | 0.33 | 0.04297 |
| GO:0008514 | organic anion transmembrane transporter ... | 57 | 3 | 0.79 | 0.04406 |
| GO:0017048 | Rho GTPase binding | 26 | 2 | 0.36 | 0.0497 |
